# Enhancer-directed gene delivery for digit regeneration based on conserved epidermal factors

**DOI:** 10.64898/2025.12.01.691633

**Authors:** David A. Brown, Katja K. Koll, Erin Brush, Grant Darner, Timothy Curtis, Thomas Dvergsten, Melissa Tran, Colleen Milligan, David Wolfson, Trevor J. Gonzalez, Sydney Jeffs, Alyssa Ehrhardt, Rochelle Bitolas, Madeleine Landau, Kendall Reitz, David S. Salven, Leslie A. Slota-Burtt, Isabel Snee, Elena Singer-Freeman, Sayuri Bhatia, Jianhong Ou, Aravind Asokan, Joshua D. Currie, Kenneth D. Poss

## Abstract

Limb loss remains a significant clinical challenge, but regenerative medicine approaches such as gene therapy offer a promising strategy to trigger endogenous regeneration programs. Optimal vector configurations and molecular targets for appendicular skeletal repair are not well defined. Here, we leveraged insights from species with a high endogenous capacity for appendage regeneration to design an enhancer-directed gene delivery platform that functions during mouse digit regeneration, a well characterized model for partial limb regeneration in mammals. Single-cell RNA sequencing of zebrafish caudal fin regeneration, combined with expression data in regenerating salamander limbs and mouse digit tips, implicated the SP family of transcription factors as conserved, epidermally-expressed mediators of appendage regrowth. Null mutants of *Sp8* demonstrated impaired limb regeneration in salamanders, while conditional knockout of *Sp6* and/or *Sp8* in the mouse basal epidermis resulted in defective bony digit tip regeneration, involving an IL-17 mediated osteoclastogenic program. Spatiotemporally focused expression of FGF8, a known target of SP factors, using a zebrafish-derived tissue regeneration enhancer element via adeno-associated viral vectors, could partially rescue digit tip regeneration in SP knockout mice and accelerate digit regeneration in wildtype mice. Our results demonstrate a contextual gene therapy approach to address limb loss based on genes like SP transcription factors conserved across multiple contexts of appendage regeneration.

**Significance Statement:** Instructing regeneration of complex structures in mammals remains an unsolved problem. Gene therapy offers a compelling approach to foster endogenous regeneration by delivering therapeutic gene products to specific cells post injury. We identified a conserved regeneration-linked epidermal transcriptional program in mouse digit regeneration centered on the SP6 and SP8 transcription factors, involving inflammatory responses from osteoclasts. We engineered AAVs harboring a zebrafish tissue regeneration enhancer to direct FGF8 expression in the epidermis after amputation. This enhancer directed delivery partially rescued impaired digit regeneration in *Sp6* and *Sp8* conditional knockout mice and accelerated regrowth in wildtype digits. Our work links developmental signaling to adult regeneration and establishes a modular, injury site specific gene therapy framework that enables new interventions for limb healing.

## Introduction

Limb loss is a major contributor to human morbidity and mortality. As of 2017, over 57 million people were living with limb loss worldwide, predominantly due to trauma, cancer, vascular disease, and congenital anomalies(1). While prosthetic technologies have advanced considerably in the last decades, they are unable to replicate the complex sensorimotor functions of whole limbs, in addition to suffering from challenges related to tissue interfacing and chronic pain. As a large, highly complex structure involving multiple tissue types, recreating a human limb stands as a considerable aspiration. Insights from animals capable of appendage regeneration have the potential to inform strategies to augment clinically relevant elements of human limb healing and tissue regeneration, such as re-epithelialization and bone regrowth.

In two well-characterized models of appendage reformation, zebrafish fins and axolotl limbs, a specialized epidermis forms over the amputation stump shortly after injury and is known as the wound epidermis, apical epithelial cap, or regeneration epidermis (RE). Observations across multiple, evolutionarily divergent species capable of limb or appendage regeneration indicate that the RE is critical for induction of a mesenchymal blastema, which contains the progenitor cells and positional information required for structural regeneration. The RE likely functions as a signaling center akin to the developing limb bud’s apical epidermal ridge (AER), with overlapping profiles of gene expression between both contexts [see Aztekin(2) for review].

During mouse digit tip regeneration, a distinct RE forms, though its formation is delayed until histolysis of the bone stump is complete (3, 4). The nail organ appears to have a critical role in mobilizing epidermal cells to reconstitute the RE after amputation, in which keratin14-positive (K14^+^) basal epidermal cells migrate from the proximal nail matrix to the amputation site in a Wnt signaling-dependent manner. Conditional epidermal deletion of β-catenin, a Wnt signaling mediator, impairs formation of the RE as well as bone regeneration(5). These findings suggest that K14^+^ basal epidermal cells of the nail matrix migrate to the regeneration epidermis and orchestrate phalangeal bone regeneration following digit amputation.

To identify and manipulate signaling pathways and associated signals from the RE that enable bony regeneration in the digit tip, we designed a transgenic strategy to enhance limb regeneration. A discovery-based approach using single-cell RNA sequencing (scRNA-seq) of the zebrafish fin RE in conjunction with comparative genomics highlighted the SP family of transcription factors as key genes in this process. Expression and knockout studies in regenerating zebrafish fins, axolotl limbs, and mouse digit tips implicated homologs of *Sp6* and *Sp8* – known developmental Wnt target genes and upstream mediators of Fgf signaling and other signaling pathways. Using a recombinant adeno-associated virus (AAV) variant known to effectively transduce limb tissue, we tested the ability of *Fgf8* gene transfer to rescue conditional knockout phenotypes and enhance wildtype mouse digit tip regeneration. Our results introduce methodology for targeted limb regeneration gene therapy based on conserved, regeneration-linked factors of the epidermis.

## Results

### Epidermal SP transcription factors are conserved mediators of appendage regeneration

To identify important factors expressed in the zebrafish caudal fin RE, we first examined gene expression profiles of basal keratinocytes during the first four days after amputation, corresponding to the timeframe of RE maturation and blastema development. For these experiments, a dual transgenic reporter line was bred, consisting of nuclear-localized mCherry in basal epidermal cells [*Tg(krt1 19e:h2az2a mCherry)*](6) and membrane-bound GFP in superficial epidermal cells [*Tg(krt4:LY EGFP)]*(7) (Fig. 1*A*). Live imaging of anesthetized fish after fin amputation demonstrated stratified expression of the transgenic reporters, with mobilization of basal cells occurring within hours of injury (*SI Appendix,* Movie S1). At each time point, dissociated cells from fin regenerates were enriched for basal epidermal cells by FACS based on transgene fluorescence followed by scRNA-seq of the sorted *krt1-19e*^+^*krt4*^-^ population (Fig 1*A*). In the integrated dataset of 17,449 cells, basal-enriched cells comprised 10.3% of the final sequenced population. Unbiased clustering of the integrated dataset resolved 20 clusters (C0–C19) encompassing basal/intermediate/superficial epithelium, mesenchyme, osteogenic lineage cells, endothelium, and immune cell populations (Fig. 1*B*; SI *Appendix*, Fig. S1*A*). Cluster identities were assigned using the most differentially expressed genes, based on literature references and canonical markers (SI *Appendix*, Dataset S1). Non-epithelial clusters likely reflect minor carryover from adjacent tissues during dissection rather than genuine inclusion in the sorted epithelial population.

**Figure 1.**
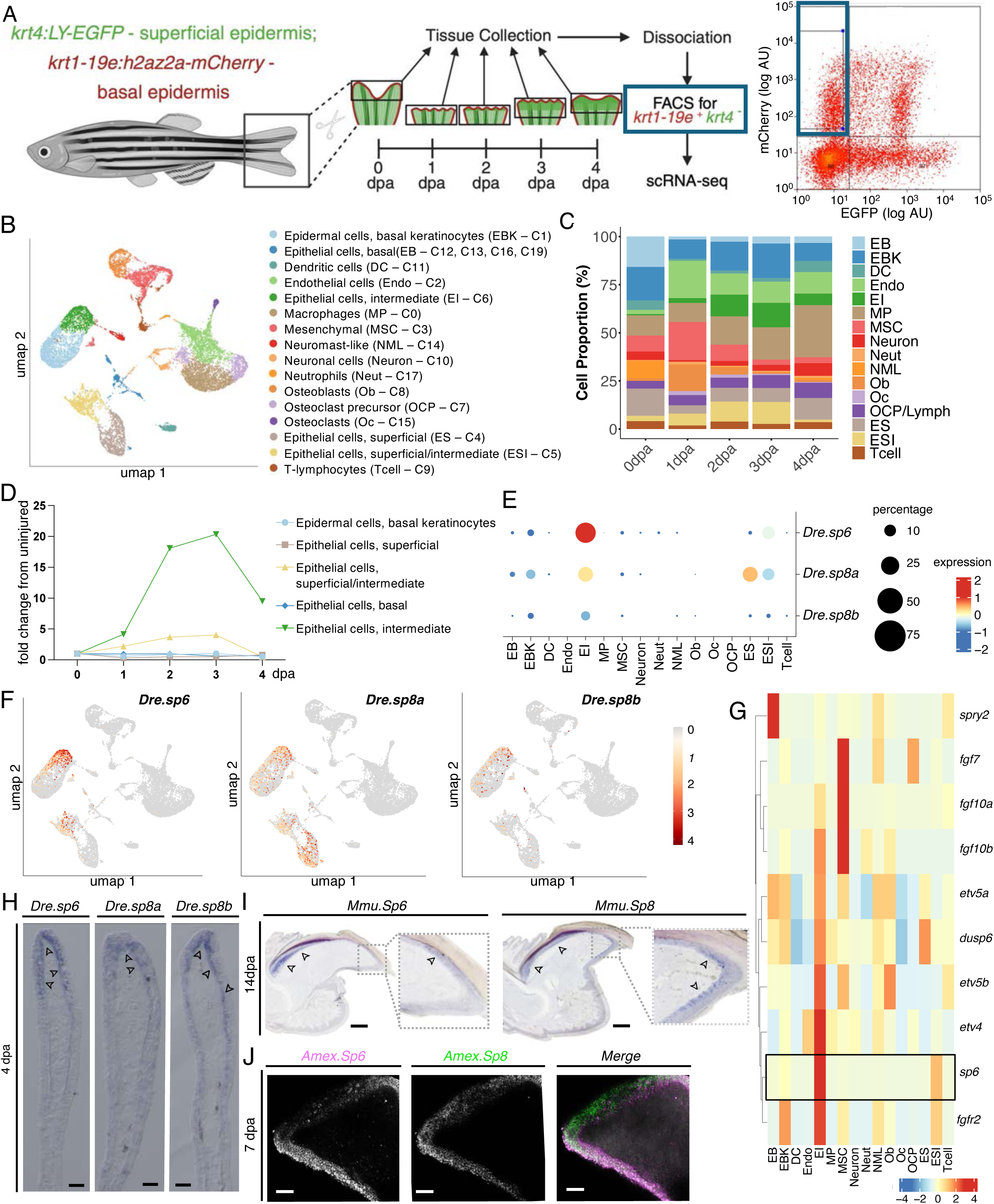
Epidermal SP transcription factors are conserved mediators of appendage regeneration. (A) Schematic of sample collection and enzymatic dissociation for RNA-seq profiling of mCherry/EGFP zebrafish fins at 0, 1, 2, 3, and 4 days post-amputation (dpa). For 0 dpa, 1.5 mm fin tissue segments were collected from the central region of intact fins. Regenerating samples were collected at 1–4 dpa from newly regenerating fin tissue. At each time point, dissociated cells were enriched for basal epidermal cells by FACS using endogenous reporter fluorescence, followed by single-cell RNA sequencing of the *krt1 19e:h2az2a mCherry^+^, krt4:LY EGFP*-population. (B) UMAP visualization of integrated scRNA-seq datasets showing cell clusters (C) with annotated identities. Assigned cell types include: epidermal cells, basal keratinocytes (EBK, C1), epithelial cells, basal (EB, C12, C13, C16, C19), dendritic cells (DC, C11), endothelial cells (Endo, C2), epithelial cells, intermediate (EI, C6), macrophages (MP, C0), mesenchymal cells (MSC, C3), neuromast-like cells (NML, C14), neurons (C10), neutrophils (Neut, C17), osteoblasts (Ob, C8), osteoclast precursors (OCP, C7), osteoclasts (Oc, C15), epithelial cells, superficial (SE, C4), epithelial cells, superficial/intermediate (ESI, C5), and T cells (C9). (C) Bar plot showing the percentage distribution of the cell types identified in (B), grouped by collection stage, illustrating changes in cellular composition across regeneration. (D) Dynamics of epithelial clusters over time (1–4 dpa), shown as fold change relative to uninjured tissue, highlighting temporal shifts in epithelial subpopulations during fin regeneration. (E) Dot plot showing differential expression of *Dre.sp6, Dre.sp8a*, and *Dre.sp8b* across the cell type clusters, illustrating their enrichment patterns. Color intensity indicates expression level (dark blue = low, red = high), while circle size represents the percentage of cells expressing each gene (small = low, large = high). (F) UMAP feature plots showing the distribution of *Dre.sp6, Dre.sp8a*, and *Dre.sp8b* expression across cell clusters. Expression levels are indicated by color intensity, with grey representing low and red representing high expression. (G) Heatmap showing expression of *Dre.sp6* and Fgf signaling pathway genes across cell types. Co-expression patterns are evident, with many Fgf pathway genes enriched in intermediate epithelial (EI) cells alongside *Dre.sp6*. (H) In situ hybridization (ISH) detecting *Dre.sp6, Dre.sp8a, and Dre.sp8b* expression in regenerating fins at 1, 4, and 7 dpa. Arrows indicate sites of transcript enrichment within the regenerating epidermis. Scale bars, 50 μm. (I) ISH for *MMu.Sp6* and *Mmu.Sp8* on sagittal sections of regenerating mouse digit tips (P3) at 14 dpa. Black arrowheads indicate domains of transcript enrichment within the regenerating nail matrix and regeneration epidermis. Scale bars, 100 μm. (J) Hybridization chain reaction whole-mount fluorescent in situ hybridization (HCR WMFISH) of *Amex.Sp6* and *Amex.Sp8* in lower limb (mid radius/ulna) at 7 days post amputation (dpa) with DAPI merge (right panel). Images represent maximum intensity projection of 25µm z-depth. Scale bars, 100 μm.

Because the RE transitions within the first 2-3 days post amputation (dpa) from a simple wound cover to an organizing epithelium that directs blastema proliferation and patterned outgrowth (8, 9), we examined temporal shifts in cell-type composition. Cell-type proportions changed over time (Fig. 1C; SI *Appendix*, Fig. S1*B–D*), with intermediate epithelium (EI) increasing most prominently at 2–3 dpa relative to uninjured fins (Fig. 1*D*). Gene ontology (GO) analysis of intermediate epithelia (EI) markers highlighted programs in ECM organization, fin regeneration, and skeletal/vascular development (SI *Appendix*, Fig. S1*E–F*), indicating that the EI functions not as a passive wound cover but as a signaling hub coordinating mesenchymal proliferation and tissue remodeling, specifically at 2–3 dpa. Within this context, among the top 30 EI-enriched genes, *Dre.sp6* emerged as a lead candidate for investigation, peaking at 2–3 dpa (SI *Appendix*, Fig. S1*G*). Across the atlas, *Dre.sp6* and *Dre.sp8* paralogs (*Dre.sp8a, Dre.sp8b*) were selectively enriched in epithelial compartments—most prominently EI at 2–3 dpa (Fig. 1*E,F*; SI *Appendix*, Fig. S1*G*). Pathway-level analyses of epithelial regeneration modules (Wnt/β-catenin, BMP/TGFβ, Hedgehog, and FGF) showed co-enrichment of *Dre.sp6* with multiple Fgf-pathway genes within IE (Fig. 1G; SI *Appendix*, Fig. S1*H–J*). This pattern echoes developmental roles of SP factors in the mammalian AER, where SP6/SP8 supports *Fgf8* expression downstream of Wnt/β-catenin(10).

Members of the SP transcription factor family are established regulators of appendage development in zebrafish fins(11) and mammalian limbs(12, 13). Consistent with this, we similarly detected Mmu SP6 and Mmu SP8 in the AER of E12.5 mouse limb buds (SI *Appendix*, Fig. S2*A*). Extending these findings to urodeles, recent work implicated *Amex.Sp6* and *Amex.Sp8* in axolotl limb development(14), and we likewise observed robust *Amex.Sp6* and *Amex.Sp8* expression in axolotl limb buds at early (stages 44–45) and mid (stages 45–46) phases (SI *Appendix*, Fig. S2*B*). Phylogeny and sequence alignments show strong conservation of the C-terminal C2H2 zinc-finger DNA-binding array with more divergent N termini across axolotl, zebrafish, mouse, and human. Mouse clusters are closest to human and predicted 3D structures are highly similar (SI Appendix, Fig S3*A,B*). Together, these data support a conserved epidermal role for the SP family in appendage development.

To test whether the regeneration-specific epidermal SP program is conserved across species and injury contexts, we examined expression in regenerating adult zebrafish, axolotl, and mouse appendages. In situ hybridization (ISH) confirmed basal/intermediate epidermal localization of *Dre.sp6*, *Dre.sp8a*, and *Dre.sp8b* in zebrafish fins at 1, 4, and 7 dpa, with the highest visible signal at 4 dpa in the distal RE (Fig. 1H; SI *Appendix*, Fig. S4*A*). In adult mouse digits, *Mmu.Sp6* and *Mmu.Sp8* were low at baseline but visible after P3 amputation, localizing to the nail bed and RE overlying the blastema area. Signals became apparent by 7 dpa, peaked at 10–12 dpa, and declined slightly by 14 dpa as differentiation progressed (SI *Appendix*, Fig. S4B; Fig. 1I), with IF substantiating nuclear Mmu SP6 and Mmu SP8 in the RE (SI *Appendix*, Fig. S4C). Critically, this induction was regeneration-specific; following P2 amputations that fail to regenerate, *MMu.Sp6* and *MMu.Sp8* remained undetectable from 7–14 dpa (SI *Appendix*, Fig. S4D). Finally, hybridization chain reaction whole-mount fluorescent in situ hybridization (HCR WMFISH) in regenerating axolotl lower limbs (mid radius/ulna) demonstrated conserved epithelial expression of *Amex.Sp6* and *Amex.Sp8* at 7, 10, and 14 dpa, with a peak at 10 dpa (Fig. 1J; SI *Appendix*, Fig. S4*E*).

Taken together, the epidermal-based *Dre.sp6* and *Dre.sp8* expression in zebrafish, its co-occurrence with epithelial pathway signatures (including Fgf), and regeneration-specific induction in axolotl limb and mouse digit tip epithelia collectively implicate SP6/SP8 as conserved epithelial mediators of appendage regeneration.

### *Sp6* and *Sp8* are required for axolotl limb and mouse digit tip regeneration

As a genetic test of the necessity of SP6 and SP8 to appendage regeneration, we generated loss-of-function mutants in zebrafish, mice, and axolotl. Zebrafish knockouts of *Dre.sp6*, *Dre.sp8a*, or *Dre.sp8b* did not result in detectable phenotypes of impaired fin regeneration (SI *Appendix*, Fig. S5*A-B*). We speculated that there exists considerable functional redundancy in the zebrafish epidermal SP family, similar to mouse SP homologs during limb development(15), so we bred double-deletion combinations of the lines. While *Dre.sp8a*; *Dre.sp8b* deletions resulted in death prior to maturity (SI *Appendix*, Fig. S5*C*), we were unable to detect abnormalities in fin regeneration in other double-deletion mutants (SI *Appendix*, Fig. S5*D*).

To test the necessity of *Amex.Sp8* in axolotl limb regeneration, we generated CRISPR–Cas9 mutants using six non-overlapping guide RNAs (gRNAs) spanning the *Amex.Sp8* coding region (Fig. 2*A*, SI *Appendix*, Fig. S6*B,C*). Unexpectedly, both F0 *Amex.Sp8* crispants and germline F1 *Amex.Sp8* knockouts exhibited no limb development abnormalities, unlike previously reported developmental phenotypes for *Mmu.Sp8* knockout mice(15). However, amputating *Amex.Sp8* KO limbs at the mid-zeugopod (radius/ulna) level resulted in a failure to correctly pattern distal skeletal elements (Fig. 2*C*). Secondary regenerates produced from tissue harvesting exhibited further degradation of skeletal patterning (Fig. 2D-E). Across independent gRNAs, approximately 26–34% of animals exhibited regeneration defects compared to regeneration in *Tyrosinase* (*Amex.Tyr*) gRNA control limbs (Fig. 2*B*). Additionally, we noted that skeletal integration between existing radius/ulna and regenerated cartilage were mismatched and showed incomplete histolysis in the majority of *Amex.Sp8* knockout regenerates (Fig 2*C*). These data demonstrate that *Amex.Sp8* is required for axolotl limb regeneration, consistent with its expression in the regeneration epidermis and its established AER-related role in promoting epithelial–mesenchymal signaling during limb outgrowth(11).

**Figure 2.**
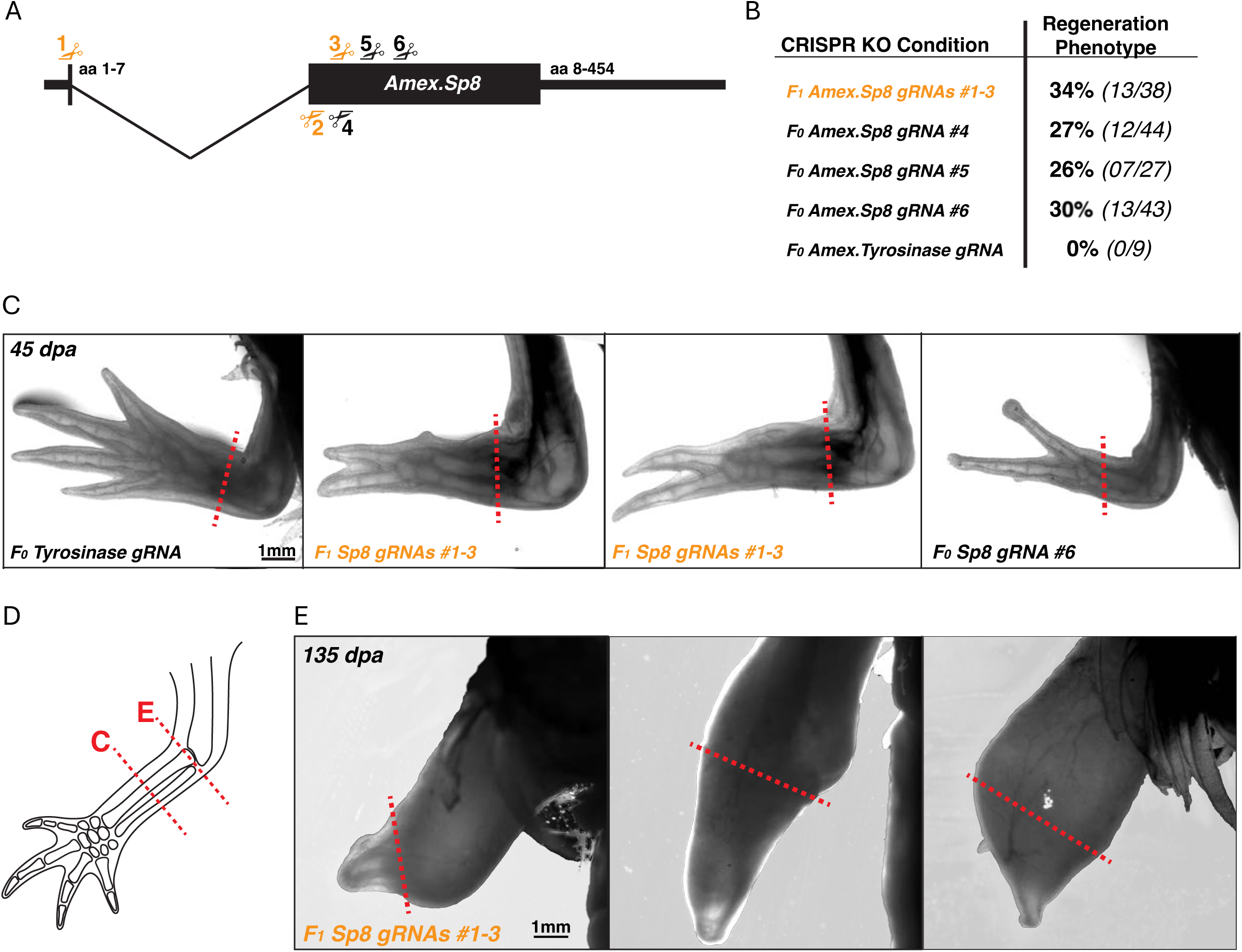
*Amex.Sp8* is dispensable for axolotl limb development, but required for limb regeneration. (A) Schematic of Amex.Sp8 genomic locus with 5’/3’ UTRs, 2 exons, and intron. Six CRISPR/Cas guide RNAs (gRNAs) are shown relative to their target on 5’ coding (top) or 3’ non-coding (bottom) strand. *Amex.Sp8* gRNAs 1-3, labeled in gold, were injected together in axolotl embryos and raised to F1 hemizygous larvae. (B) Table summarizing the proportion of axolotls with regeneration-specific limb mis-patterning. All axolotls displayed normally patterned limbs prior to amputation at mid-radius/ulna level. Age matched F0 CRISPR/Cas *Tyrosinase* (*Amex.Tyr*) knockout axolotls were used as a negative control for proper regeneration. (C) Representative images of limb regenerates in control F0 *Amex.Tyr* CRISPR KO animals (left panel) and F1 and F0 *Amex.Sp8* KO larvae axolotls at 35 dpa. Amputation planes are indicated by red dashed lines. Scale bar, 1 mm. (D) Cartoon representation of primary amputation planes represented in (C), and secondary amputations with resulting regenerates reported in (E). (E) Secondary regenerates from F1 *Amex.Sp8* animals amputated just distal to the elbow. Secondary amputations for tissue harvesting were performed approximately 115 days after the primary amputations in (C). Images were taken 135 days post-secondary amputation. Scale bar, 1 mm.

We also generated basal epidermis-specific inducible knockout mouse lines harboring homozygous *Krt14:CreER* alleles and homozygous floxed *Mmu.Sp6* and/or *Mmu.Sp8* alleles (*K14:CreER^+/+^;Sp6^fl/fl^, K14:CreER^+/+^;Sp8^fl/fl^,* and *K14:CreER^+/+^;Sp6^fl/fl^*;*Sp8^fl/fl^*). This allowed loss-of-function studies in developmentally normal adults, which would otherwise have an array of limb defects in conventional germline knockouts(15). To test the contribution of SP factors to digit regeneration in the epidermis specifically, *Krt14:CreER*-mediated recombination was induced prior to digit tip (P3) amputation (Fig. 3*A*). Although we detected baseline expression (leakiness) of the uninduced *Krt14:CreER* transgene (SI *Appendix*, Fig. S6*A*), conditional KO (cKO) mice exhibited a significant impairment in P3 regeneration in comparison with tamoxifen-treated WT controls. Histologically, we noted a delay in formation of the distal epidermal covering at 10 dpa (Fig. 3*B*), abrogated blastema formation at 14 dpa, and ultimately shorter and less voluminous bone regenerate at 28 dpa (Fig. 3*B*). Masson’s trichrome staining showed smaller, less mature regenerates in *Mmu.Sp6* and *Mmu.Sp8* single cKOs versus controls, with 14 dpa distal stumps capped by abundant collagen with reduced bone formation. By 28 dpa, control digits had formed a well-structured distal phalanx, whereas cKOs retained diminished ossified regenerate and disorganized extracellular matrix, indicating attenuated osteogenesis (Fig. 3*C*).

**Figure 3.**
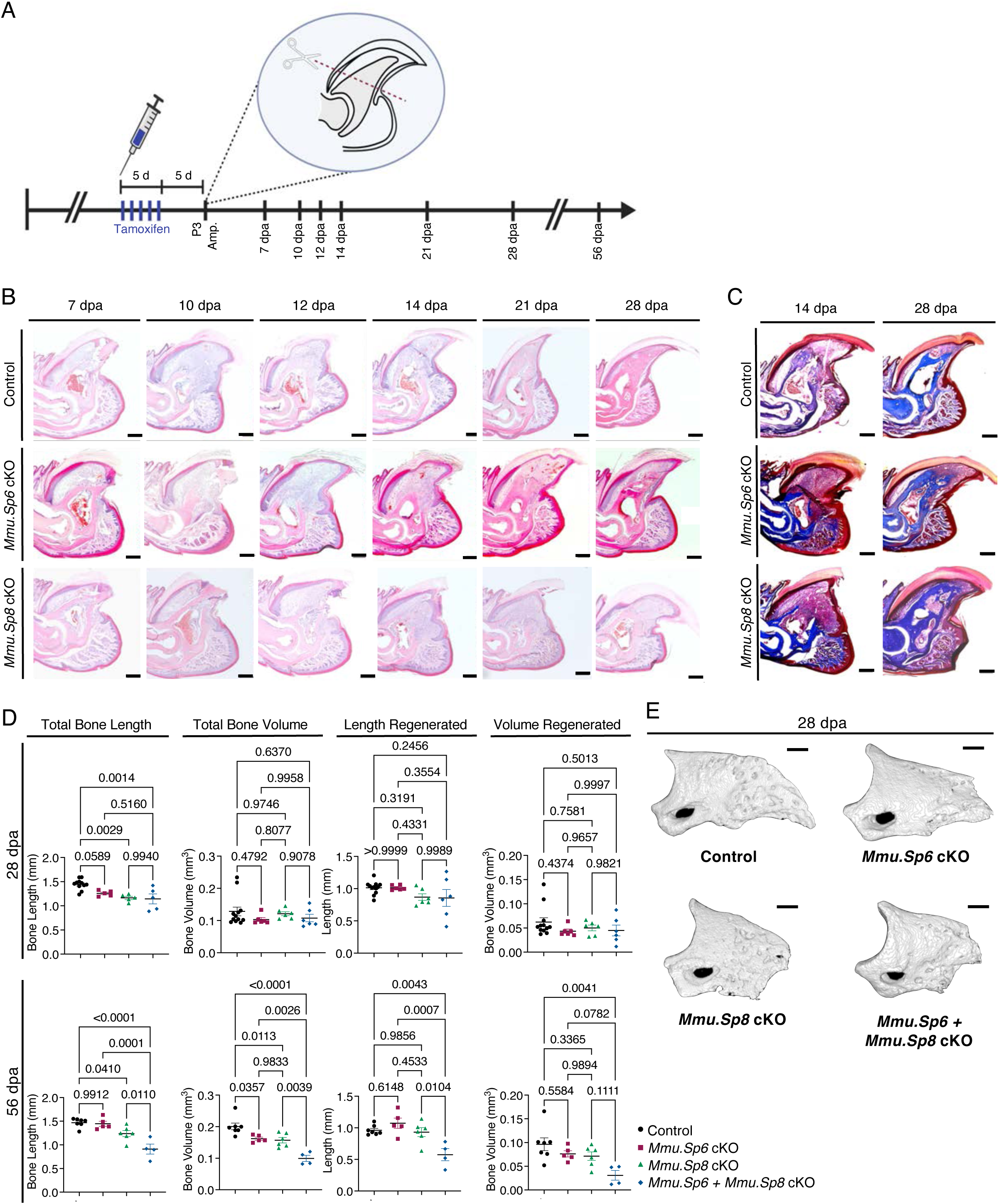
*Sp6* and *Sp8* are required for mouse digit tip regeneration. (A) Experimental design of *Mmu.Sp6, Sp8*, and *Sp6+Sp8* conditional knockout models using the mouse digit tip (P3, third phalanx) amputation (amp.) model with tamoxifen induction. The timeline indicates tissue collection at 7–56 days post-amputation (dpa). (B) H&E time course of digit regeneration following P3 amputation in *Sp6* and *Sp8* cKO mice compared with controls. Compared with controls, both cKO genotypes show delayed/defective regeneration, characterized by reduced or disorganized blastema and diminished distal bone outgrowth at indicated stages. Scale bar, 100 μm. (C) Masson’s trichrome staining of digit tips at 14 and 28 dpa in *Sp6* cKO and *Sp8* cKO mice compared with controls. Scale bar, 100 μm. (D) Quantification of digit tip regeneration from micro-CT analyses in tamoxifen-induced WT controls, *Sp6* cKO, *Sp8* cKO, and *Sp6+Sp8* cKO mice at 28 and 56 dpa. Total bone length and volume, as well as regenerated length and volume, were measured. Data are presented as mean ± SEM (at 28 dpa: Control n=11, *Mmu.Sp6* cKO *n*=5, *Mmu.Sp8* cKO *n*=5, *Sp6+Sp8* cKO: *n* = 5; at 56 dpa: Control n=7, *Mmu.Sp6* cKO *n*=5, *Mmu.Sp8* cKO *n*=4, *Sp6+Sp8* cKO: *n* = 4) (E) Representative 3D micro-CT reconstructions of digit tip regeneration in control, *Sp6* cKO, *Sp8* cKO, and *Sp6+Sp8* cKO mice at 28 dpa. Scale bar, 200 μm.

Among cKOs, quantitative micro-CT confirmed significant reductions in total bone length (13% for *Mmu.Sp6*, 19% for *Mmu.Sp8*, 21% for *Mmu.Sp6*+*Mmu.Sp8*) at 28 dpa. Similar to observations during limb development(15), *Mmu.Sp6* cKO produced the mildest defects, followed by *Mmu.Sp8* and then the combination *Mmu.Sp6*+*Mmu.Sp8* cKO. *Mmu.Sp6+Mmu.Sp8* cKO nearly abolished regrowth at 28 and 56 dpa: instead of a regenerated bone, only a blunt ossified stump or minimal new bone formed beyond the amputation plane (Fig. 3*D,E*). Longer-term evaluation of bone at 56 dpa - approximately double the amount of time required for complete regeneration - suggests that the regeneration phenotype is not only a delay in the process but a persistent impairment. Mice with *Mmu.Sp6*+*Mmu.Sp8* cKOs exhibited reductions in all measured parameters at 56 dpa: total bone length (28%), total bone volume (50%), regenerate bone length (40%), and regenerate bone volume (68%). Notably, nail growth was also delayed in *Mmu.Sp8* cKOs at 28 and 56 dpa (SI *Appendix*, Fig. 7*A*). Interestingly, we detected higher rates of cell cycling in multiple anatomic regions of all SP cKO mutants (SI *Appendix*, Fig. S8*A,B*).

### IL-17-mediated osteoclast signaling pathway is disrupted in Sp6 and Sp8 mutant digit tip regenerates

To explore mechanisms by which *Mmu.Sp6* and *Mmu.Sp8* regulate mouse digit tip regeneration, we performed bulk RNA-seq analysis of regenerating digit tissues in cKO mice versus tamoxifen-injected wildtype controls. At 14 dpa, corresponding to the peak blastema stage during adult mouse digit regeneration, RNA-seq from *Mmu.Sp6* cKO, *Mmu.Sp8* cKO and *Mmu.Sp6+Mmu.Sp8* cKO mice confirmed reductions in *Mmu.Sp6* expression to 3.9% and *Mmu.Sp8* expression to 24% of wildtype controls in the respective cKO lines (SI *Appendix*, Fig S9*A*). Mutant mice exhibited a pronounced inflammatory transcriptomic signature with unique differentially expressed genes (DEGs). Pathway analysis of genes elevated in *Mmu.Sp6+Mmu.Sp8* cKOs showed strong enrichment for inflammatory remodeling, with IL-17 signaling among the most significantly affected pathways, together with associated categories such as T-helper-17 differentiation, cytokine activity, and osteoclast differentiation (SI *Appendix*, Fig. S9*B,C,D*). Conversely, epidermal differentiation programs were relatively diminished (SI *Appendix* Fig. S9*B,C*). Further analysis of DEGs identified a shared set of IL-17–axis (*Il17a*, *Cxcl5*, *Cxcl13*) and osteoclast-related genes (*Mmp9*, *Oscar, Il1b*) that were upregulated in all SP cKO lines compared to controls (SI *Appendix*, Fig. S9*E,F,G*). The transcriptional induction of *Il17a* was accompanied by a significantly increased expression of *Rorc* which encodes RORγt, the key transcription factor of Th17 cells, in all mutants (SI *Appendix*, Fig. S9*E,F,G*). Th17 cells (characterized by RORγt expression) are a distinct lineage of CD4⁺ T helper cells that produce IL-17A and drive proinflammatory responses(16). IL-17 signaling is implicated in chronic inflammation and in promoting osteoclast-mediated bone resorption. Consistent with this, IL-17 has been shown to facilitate osteoclast differentiation and pathological bone loss *in vivo*(17). Thus, the observed upregulation of *Il17a* (along with *Il1b* and chemotactic factors) in *Mmu.Sp6* and *Mmu.Sp8*-deficient digits suggests downstream inflammatory osteoclastic activity.

As an alternative test for transcriptional osteoclast activation in SP mutants via RNAseq, we assessed osteoclast activity histologically. Immunofluorescence revealed increased Cathepsin K⁺ cells within the blastema of all mutant genotypes relative to controls, with the most pronounced increase in *Mmu.Sp6* cKO mice (SI *Appendix*, Fig. S9*H,I*). Quantification of Cathepsin K⁺ cells per blastema area showed a significant elevation in *Mmu.Sp6* cKO mice vs. controls (*p* = 0.0034), with trends in *Mmu.Sp8* cKO and *Mmu.Sp6+Mmu.Sp8* cKO animals (SI *Appendix*, Fig. S9*H*). These histological data are concordant with the RNA-seq signature indicating enhanced osteoclast activation in the mutants (SI *Appendix*, Fig. S9*E,F,G*).

Finally, to explore potential gene regulatory mechanisms controlling IL-17-mediated osteoclast activation, we performed *in silico* motif analysis. We detected GC-rich consensus motifs for SP family members (*Sp1*–*Sp4*) within promoters of *Rorc* and *Il17a* (SI *Appendix*, Fig. S10*A*) and mapped multiple predicted binding sites across these promoters (SI *Appendix*, Fig. S10*B*). Notably, independent work showed that SP3 can suppress RORγt expression and prevent Th17 cell differentiation(17, 18). Our motif analysis therefore implicates *Rorc* and *Il17a* as negatively-regulated targets of SP family regulators during digit tip regeneration.

### Targeted *Fgf8* gene expression in regenerating digit tips enhances bone regrowth

Previous work has demonstrated that systemic introduction of AAVs in mice can spatiotemporally direct cargo expression to a myocardial or spinal cord injury when engineered with a zebrafish tissue regeneration enhancer element (TREE)(19, 20). TREEs are identifiable by regeneration-specific chromatin accessibility, an approach which has uncovered similar classes of regulatory elements in *Drosophila* imaginal wing discs(21–23), acoel worms(24), mouse bone(25), and zebrafish hearts(26, 27). For this study, we utilized the zebrafish-derived *leptin b*-linked enhancer (LEN), which was previously shown to direct gene expression in injured transgenic mouse hearts and digits(20), and in injured hearts of AAV-transduced mice.

To test whether this strategy could be applied to adult regenerating digit tips, we first screened AAV serotypes carrying the ubiquitous expression cassette *CBh:GFP*, which allowed us to identify transduced cell types in the digit tip. AAVs were administered by retroorbital injection at three time points after amputation, followed by digit harvest at 14 dpa and assessment of transduction by histology. Both AAV6 and AAV8 exhibited transduction of the blastema when injected at 10 dpa, though little signal was seen in the distal epidermis (SI *Appendix*, Fig. S11*A*). We then tested the engineered capsid variant AAV.cc47, which was previously developed by iterative cycling of AAV9 libraries in pigs, mice, and non-human primates, followed by testing of gene transfer efficiencies in mice(28). Transduction of the distal epidermis was observed with AAV.cc47 when injected at 7 dpa and 10 dpa, demonstrating that it is applicable for gene transfer to this anatomic region as well (SI *Appendix*, Fig. S11*A*). In liver sections, injected samples exhibited robust, mosaic GFP fluorescence surrounding DAPI-labeled nuclei, whereas uninjected controls showed no detectable GFP signal, indicating effective systemic delivery (SI *Appendix*, Fig. S11*B*).

As a first test of the ability of TREEs to confine gene expression to the regenerating digit, we subcloned *LEN* upstream of the *Hsp68* permissive promoter followed by the GFP or luciferase coding sequence into an AAV genomes. These genomes were then packaged into the AAV.cc47 vector, followed by retroorbital injection at 10 dpa (Fig. 4*A*), which avoids vector genome dilution during the initial histolysis period and allows maximum transgene expression at the peak of the blastema (14 dpa). *LEN*-driven luciferase expression was sustained at the digit tip amputation site after a single dose of AAV, which persisted over approximately 10 days (Fig. 4*B*). Remarkably, even after systemic delivery, detectable expression was absent from all other tissues except the liver, which is the primary site of AAV clearance. Histology indicated transient expression directed by *LEN* in mesenchyme and the distal epidermis (Fig. 4C). These experiments suggest that the AAV TREE system is capable of injury site-specific delivery of genetic cargos to distal digit tip tissues.

**Figure 4.**
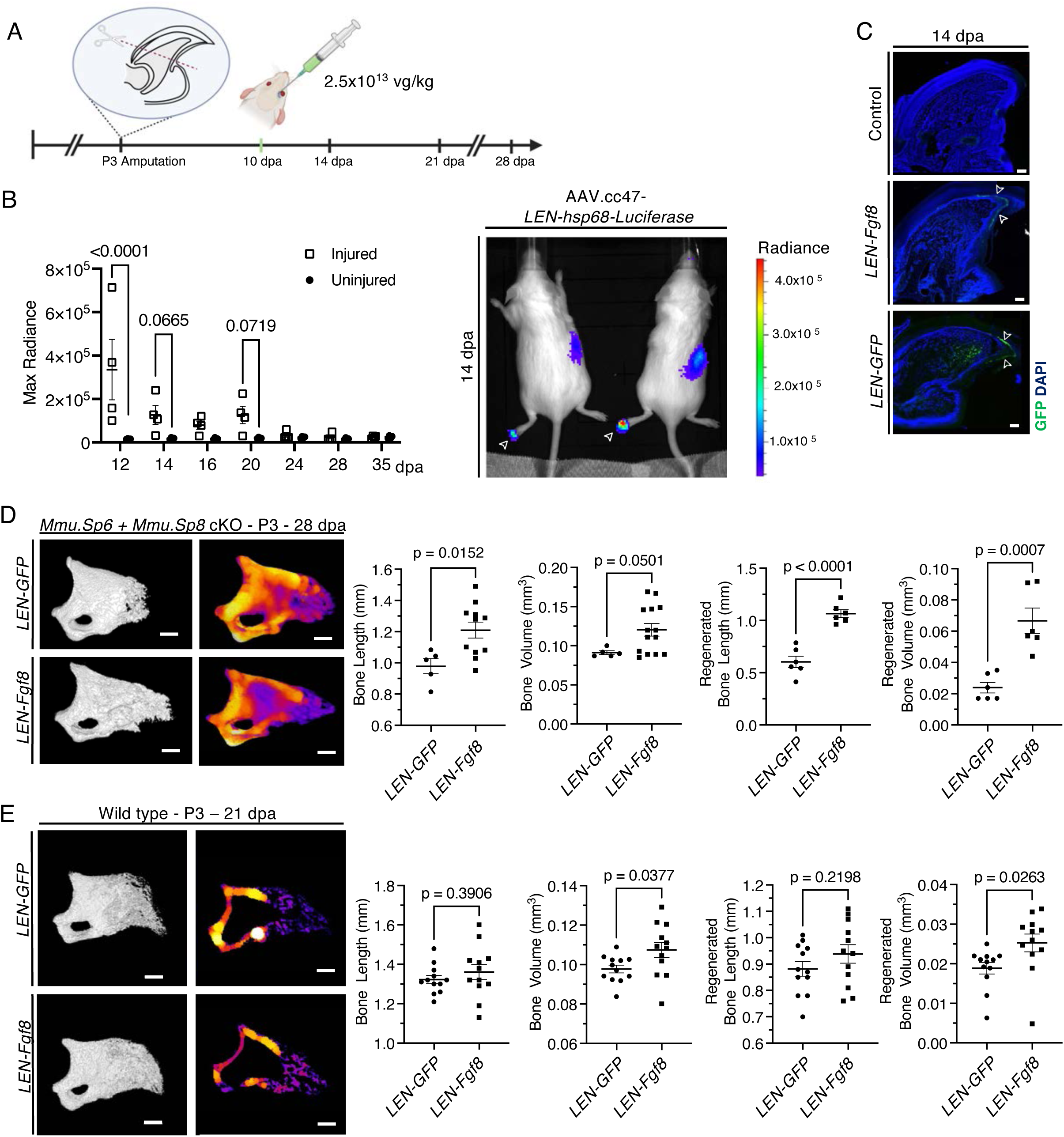
Targeted *Fgf8* gene transfer to the digit regeneration epidermis enhances bone regrowth. (A) AAV was systemically administered to 2–6-month-old mice via retro-orbital injection at a dose of 2.5 × 10¹³ viral genomes per kilogram of body weight. The injection was performed 10 days after P3 amputation (10 dpa), and tissues were collected at 14, 21, and 28 dpa. (B) Bioluminescence imaging of regenerating digits (P3) following AAV.cc47-*LEN-hsp68-Luciferase* delivery. Representative images at 14 dpa (top) show luciferase activity localized to the injured digits (arrow). Quantification of radiance (bottom) is shown at 12, 14, 16, 20, 24, 28, and 35 dpa in injured versus uninjured digits. Data are presented as mean ± SEM (*n* = 4); *p*-values are indicated. (C) GFP expression in regenerating P3 digits at 14 dpa following injection of AAV.cc47-*LEN-hsp68-GFP* (*LEN-GFP*) or AAV.cc47-*LEN-hsp68-FGF8-P2A-GFP* (*LEN-Fgf8*), compared with control. GFP (green) marks transgene expression, and nuclei are counterstained with DAPI (blue). Arrows indicate GFP expression in the distal epidermis. Scale bars, 100 μm. (D) Micro-CT reconstructions and quantitative analysis of digit regeneration in *Sp6+Sp8* cKO mice at 28 dpa following injection of *LEN-GFP* or *LEN-FGF8*. Representative 3D reconstructions (left) and heat maps of mineralized tissue (middle) show bone morphology and density. Quantification (right) includes total bone length, bone volume, regenerated bone length, and regenerated bone volume. Data are presented as mean ± SEM (*n* = 6 per group); *p*-values are indicated. Scale bars, 200 μm. (E) Micro-CT reconstructions and quantitative analysis of digit regeneration in wild-type mice (P3) at 21 dpa following injection of *LEN-GFP* or *LEN-Fgf8* vectors. Representative 3D reconstructions (left) and heat maps of mineralized tissue (middle) show bone morphology and density. Quantification (right) includes total bone length, bone volume, regenerated bone length, and regenerated bone volume. Data are presented as mean ± SEM (n = 12 per group); p-values are indicated. Scale bars, 200 μm.

To attempt to restore digit regeneration in SP mutants using AAV.cc47 and *LEN*, we selected a secreted cargo based on developmental contexts demonstrating a tight functional linkage between epidermal SP factors and FGF8 signaling. In chick limb buds, forced *Sp8* expression broadens the *Fgf8* domain, whereas *Sp8* knockdown contracts it, producing a hypoplastic AER and distal skeletal truncations(11). In mice, germline loss of *Mmu.Sp8* abolishes *Fgf8* induction in the AER(15) and *Mmu.Sp6* loss-of-function similarly shortens the *Fgf8* expression field during limb outgrowth(29). Recent ChIP-seq and RNA-seq datasets from embryonic limb ectoderm further suggest that Mmu SP8 regulates *Fgf8* both by direct binding to GC-rich SP consensus motifs and indirectly through cooperative interactions with Dlx5. These datasets identify additional AER genes under Mmu SP8 control, including En1, Wnt7a, Rspo2, and even Mmu SP6, supporting a broader role for SP8 in orchestrating epidermal signaling centers(10).

To examine whether TREE-directed FGF8 could affect regeneration in *Mmu.Sp6+Mmu.Sp8* cKO digit tips, we packaged an AAV genome containing LEN–Hsp68–Fgf8–P2A–GFP (hereafter LEN– Fgf8) in the AAV.cc47 capsid variant. When LEN-Fgf8 was administered to *Mmu.Sp6+Mmu.Sp8* cKO mice, GFP expression from the bicistronic cassette was confined to the distal epidermis at 14 dpa, indicating specificity of enhancer activity within the regenerating digit tip (Fig. 4*C*). By 28 dpa, treated digits showed 32% greater total bone volume, 24% greater regenerated bone length, and regenerated bone volume compared with LEN-GFP controls (Fig. 4*D*). These findings indicate that AAV-mediated Fgf8 delivery counteracts the osteolytic defects caused by *Sp6/8* deletion and promotes bony regeneration.

To test whether targeted expression of FGF8 could enhance digit regeneration in wildtype mice, we performed amputations at normally regenerative (P3) or normally non-regenerative (P2) levels of the digit, followed by systemic administration of LEN-Fgf8 at 10 dpa. After 14 dpa, we again observed distal expression of GFP after P3 amputation (Fig. 4*C*), indicating the expected specificity of the virus. Interestingly, little expression of GFP was seen in the distal RE after P2 amputations (SI *Appendix*, Fig. S11*C*), indicating a reduced ability of the virus to access tissue at this level. Micro-CT was performed on P3-amputated digits at 21 dpa, which was chosen as a midpoint of phalangeal regeneration to assess whether the construct would expedite the normal regenerative process. Here, we measured a 34% increase in bony regenerate volume (Fig. 4*E*), suggesting that super-physiologic levels of FGF8 in the distal epidermis can augment this process. We did not observe an effect on P2 regeneration, likely owing to diminished transduction of the virus at this level or the much higher barrier to regeneration for these structures (SI *Appendix*, Fig. S11*C*).

## Discussion

Here we find that a cross-species enhancer-directed AAV strategy aimed at the SP-Fgf signaling axis rescues and accelerates mouse digit regeneration, presenting an early but potentially generalizable blueprint for limb repair vectors. By identifying key cell types of the zebrafish fin RE, we uncovered a conserved epidermal module centered on *Sp6* and *Sp8*. Deletion of these factors impaired blastema formation and bone regrowth in mouse digit and axolotl limb, whereas zebrafish TREE-directed delivery of the SP downstream effector FGF8 partially rescued the knockout phenotype and accelerated healing in wildtype digits.

These results connect and extend several previous lines of investigation. SP family members have long been recognized as AER regulators during vertebrate limb development, acting upstream of Wnt and Fgf signaling to coordinate outgrowth(11, 12, 15). Yet their relevance to post-embryonic regeneration remained largely inferential, based on correlative expression in zebrafish fins and axolotl limbs(9, 14). Our loss-of-function data provide causal evidence that SP6 and SP8 activity within the RE is indispensable for bone restoration in both amphibian and mammalian contexts, supporting the idea that developmental AER programs are redeployed during adult appendage repair. Our inability to detect a fin regeneration phenotype in zebrafish may be due to the redundancy of the zebrafish genome, which underwent a genome duplication and thus has additional paralogs. Also, fin regeneration has considerable robustness, as only a small number of genetic lesions have been found to disrupt the process despite several large-scale forward genetic screens(30–32).

A potential mechanistic theme suggested by our datasets is that loss of epidermal *Mmu.Sp6* and *Mmu.Sp8* skews the regenerate toward an IL 17–dominant inflammatory state coupled to exuberant osteoclastogenesis. Because tightly timed histolysis is physiologic in MDT regeneration, these findings suggest that the RE does not only supply morphogens but also sets the tempo of inflammatory bone remodeling, which allows progression of regeneration from histolysis to osteogenesis. *Mmu.Sp6* and *Mmu.Sp8* loss appears to reset this control point toward sustained resorption that is incompatible with full blastema maturation and patterned osteogenesis.

An enhancer-directed AAV strategy adds a layer of tissue-level precision that could help lower systemic doses and thereby mitigate many of the hepatotoxic and thrombotic toxicities now seen at high vector doses(33). The progressive clinical implementation of AAV products by targeted regulatory elements (e.g. Elevidys, Roctavian, and Hemgenix) emphasizes the viability of vector platforms like the one used here(34). Because the *LEN-Hsp68* cassette fits within standard AAV payload limits, it should transfer readily into current good-manufacturing practice workflows and, in principle, could be swapped for other orthologous regulatory elements to tailor expression to different limb or digit compartments (e.g., blastema or nail matrix). Moreover, the successful use of a zebrafish enhancer to drive FGF8 in mouse digits adds additional context to the applicability of TREEs as therapeutic AAV drivers(19).

## Materials and Methods

### Nomenclature

Gene and protein nomenclature follows ZFIN (zebrafish), axolotl community, and MGI (mouse) guidelines. Gene symbols are italicized and species specific in capitalization; proteins are non italicized with capitalization per species. Where needed to disambiguate species, we use period separated prefixes that are not part of the symbol (Dre., Amex., Mmu.).

### Zebrafish experiments

Wildtype or transgenic male and female zebrafish of the outbred Ekkwill (EK) strain were used for this study. Three to twelve-month old zebrafish were used for experiments. The *Tg(krt4:LY EGFP)sq18Tg* strain (6) was kindly donated by Thomas Carney. Zebrafish strain *Tg(krt1 19e:h2az2a mCherry)pd309Tg* was generated in the Poss lab as previously described(35). A dual reporter line composed of homozygous combinations of both alleles was bred, using fluorescence of live animals for screening.

Caudal fin amputations were performed with a transverse amputation approximately 2/3 of the distance from the distal fin surface to the caudal peduncle, using razor blades. Fish were anesthetized for all procedures in 0.75% v/v 2-phenoxyethanol (Sigma-Aldrich) in fish water. At specified time points, 2 mm of the distal most tissue was collected from the fin regenerate for analysis. Because of our study focus on the RE, uninjured samples were collected from the distal most 2 mm of uninjured fin containing the terminal edge of the epithelium, as opposed to the uninjured fin regions corresponding to each time point of regeneration. All work with zebrafish was performed in accordance with Duke University IACUC guidelines.

Zebrafish deletion lines were created by CRISPR/Cas-9 genome editing. gRNA pairs were synthesized by IDT (custom ALT-R) using homologous sequences shown in SI *Appendix*, Table S1, with resulting genomic deletions of 3.0-5.4 kb. Each gRNA pair (500 pL of 30 ng/μl) was combined with 300 ng/μl Cas9 protein (PNA Bio) and 0.05% Phenol Red (Sigma-Aldrich), followed by microinjection into one-cell zebrafish embryos within 30 minutes of fertilization. Sperm samples of F_0_ males were collected after sexual maturity and genotyped. Positive animals were outcrossed with EK wildtype females. Deletions in F_1_ adults were confirmed by PCR with primer pairs listed in SI *Appendix*, Table S2, followed by Sanger sequencing (Genewiz, Azenta). Homozygous F_2_ progeny were bred by incrossing. Alleles were designated as *sp6^pd413^*, *sp8a^pd414^*, and *sp8b^pd415^*.

### Axolotl experiments

Axolotls (*Ambystoma mexicanum*) were raised and used in accordance with the Wake Forest University Institutional Animal Care and Use Committee (protocols A23-118 and A22-173). All experiments were performed in the leucistic (*d/d*) strain of axolotl. Larval axolotls ranging in size from 3-10cm snout to cloaca were anesthetized by immersion in 0.007% Benzocaine (w/v) in holding water until cessation of movement. Amputations were performed at lower forelimb level (zeugopod) bisecting radius-ulna between hand and elbow. Protruding skeletal elements were trimmed post-limb resection before returning animals to holding water for recovery. Progress of limb regeneration was assessed at regular intervals by anesthetization and imaging using a Zeiss V16 fluorescent stereoscope.

CRISPR/Cas knockout axolotls were created consistent with previously published protocols. Briefly, one-cell stage *d/d* axolotl embryos were microinjected with 5 nanoliters per embryo of a 10µl injection mix containing 1µl 10X Cas9 Buffer and pre-complexed Cas9 protein (5 µg, IDT) with single guide RNA (4 µg, Synthego). For *Amex.Sp8* sgRNA #1-3 injections, equimolar volumes of individual Cas9/sgRNA injection mixes were combined for embryo microinjection. *Amex.Tyr* sgRNA embryo microinjection was conducted in parallel to gauge embryo microinjection viability and general CRISPR/Cas knockout efficacy. Uninjected clutch mates and *Amex.Tyr* KOs were used as limb regeneration controls in F0 Crispant (mosaic) animals. gRNA sequences are listed in SI *Appendix*, Table S1. F1 Amex.Sp8 gRNA #1-3 animals have been assigned the formal allele designation tm(*Sp8^142v6D94/+^*)^Jcurr^.

Tissue from embryo tail tips or larval finger clips from presumptive CRISPR/Cas Sp8 knockouts and Tyrosinase and wildtype controls was collected from anesthetized animals and incubated in lysis buffer (10 mM Tris HCL pH 7.5, 50 mM KCl, 0.3% Tween, 1 mM EDTA) with fresh Proteinase K at 200 µg/ml). Samples were then incubated in a thermocycler at 55 °C for one hour, 95 °C for 15 minutes, and then stored at 4 °C. PCR reactions were performed with KAPA2G Fast HotStart Genotyping Mix (Roche) with a 65 °C annealing temperature and one minute elongation time to produce a wildtype band of approximately 529 bp of the *Sp8* genomic locus. PCR primers are listed in SI *Appendix*, Table S2.

### Mouse experiments

Frozen embryos carrying the conditional *Sp6^tm1Ibmm^* allele on a CD1.129 background (Infrafrontier/EMMA EM:02449; donor: Claude Szpirer, Ph.D.) were obtained through Infrafrontier/EMMA via the Monterotondo Mouse Clinic and rederived at Duke. Mice carrying floxed *Sp8^tm2Smb/J^* (JAX stock 023415; RRID: IMSR_JAX:023415), the keratin 14 CreERT driver Tg(KRT14 cre/ERT)20Efu/J (JAX 005107; RRID: IMSR_JAX:005107), and B6.129 *Gt(ROSA)26Sor^tm1(cre/ERT2)Tyj/J^* (JAX 008463; RRID: IMSR_JAX:008463) were purchased from The Jackson Laboratory. For experimental cohorts, floxed alleles were bred to homozygosity (e.g., *Sp6*^fl/fl^), and Cre drivers were maintained hemizygous; genotyping was performed on tail biopsies by PCR using primers in SI *Appendix*, Table S2. Wild type CD 1 (ICR) outbred mice were obtained from The Jackson Laboratory. Adult mice aged 2–6 months of both sexes were used. All procedures were approved by the Duke University IACUC.

For induction of conditional knockouts, mice were administered 75 mg/kg body weight of tamoxifen (Apex Bio) diluted in corn oil (Millipore Sigma) by intraperitoneal injection daily for 5 consecutive days, followed by 5 days of rest before digit amputation. For all mouse procedures, mice were anesthetized with isoflurane and oxygen in an induction chamber. Digit amputations were performed with a #10 scalpel and stereomicroscope. For distal phalangeal (P3) amputations, the distal 2/3 of P3 was removed with a transverse amputation, ensuring at least half of the nail matrix remained. For middle phalangeal (P2) amputations, a transverse cut was made at the midpoint of P2. Mice were given 5-10 mg/kg body weight of meloxicam subcutaneously twice daily for 48 hours post-procedure. Digits were harvested by sacrifice of the animal and processing of the tissue as described in subsequent sections.

While RNA-seq confirmed robust depletion of *Mmu.Sp6* and *Mmu.Sp8* transcripts in the tamoxifen-induced mutants, we did not maintain CreERT2-negative, floxed-only colonies to assess potential effects of the loxP insertions in isolation. Importantly, both the *Mmu.Sp6*^fl/fl^ and *Mmu.Sp8*^fl/fl^ alleles have been reported as phenotypically silent when homozygous in other studies(36–38). Moreover, because the loxP sites reside in intronic regions and the neo selection cassette was excised during allele construction, insertional hypomorphy is unlikely. Consistent with previous reports (39), we observed baseline (“leaky”) recombination from the uninduced K14-CreERT2 transgene, precluding the use of vehicle-treated CreER-positive animals as controls. Instead, we compared tamoxifen-treated mutants with age-matched, tamoxifen-treated wildtype mice to control for drug effects. We have not observed limb abnormalities in any stable homozygous Cre-loxP lines, suggesting that unintended Cre activity in the basal epidermis arises predominantly after limb morphogenesis.

### Histology and staining

Immunofluorescence staining of mouse digits was performed as follows. Digits were harvested and fixed in 4% paraformaldehyde at 4 °C for 24–48 h, then decalcified in Decalcifier I (Surgipath, Leica Biosystems, Richmond, IL). For paraffin processing, samples were washed in phosphate buffered saline (PBS), passed through a graded ethanol series, and sequentially immersed in xylenes and liquid paraffin using the Tissue Tek VIP Processor and Tissue Tek TEC Embedding Center (Sakura Finetek USA, Inc.). Sagittal sections (5 µm) were prepared in collaboration with the BioRepository & Precision Pathology Center at Duke University. For staining, slides were baked at 60 °C for 45–60 min followed by 15 min at 37 °C, deparaffinized in xylenes, rehydrated through graded ethanol to DI H₂O, and subjected to antigen retrieval in pH buffered 0.1% trypsin (37 °C, 15 min). Sections were permeabilized with 0.4% Triton X 100 in PBS for 10 min and blocked at room temperature for 30 min in 5% goat serum with 0.05% Triton X 100 in PBS. Primary antibodies against SP6 (Proteintech), SP8 (Novus Biologicals), Ki67 (Abcam), GFP (Thermo Fisher), and cathepsin K (Proteintech) were diluted in 0.05% Triton X 100 with 1% neonatal calf serum and applied overnight at 4 °C. After PBS washes, slides were incubated at 37 °C for 1 h with Alexa Fluor goat anti rabbit IgG secondary antibodies (594 or 488; 1:500). Following additional PBS washes, slides were mounted with Vectashield Mounting Medium containing DAPI. Imaging was performed on Zeiss LSM 700 or LSM 880 confocal microscopes. Tile scanned z stacks were stitched using ZEN software, and images were processed in ImageJ (USA).

In situ hybridization for zebrafish transcripts was performed on cryosections of fin samples fixed in 4% paraformaldehyde as described previously by Tornini et al(40). Riboprobe templates were synthesized by IDT using the sequences listed in SI *Appendix*, Dataset S2, including a 5’ SP6 promoter and a 3’ reverse-orientation T7 promoter. RNA was then produced with T7 RNA polymerase (New England Biolabs) and labeled using a DIG RNA Labeling Mix (Roche). In situ hybridization was performed using an InSituPro VSi robot (Intavis). Imaging was performed on a Leica DM6000B compound microscope.

Whole mount hybridization chain reaction fluorescent in situ hybridization (HCR FISH) v3.0 was performed according to previously published protocols (Molecular Instruments)(41). Samples (stage 44–46 axolotl larvae or harvested limb tissue) were fixed overnight at 4 °C in 4% paraformaldehyde in PBS. After three washes in PBST (PBS + 0.01% Triton X 100), tissues were dehydrated on ice through a methanol series in PBST (25%, 50%, 75%, 100% MeOH; 10 min each) and stored at −20 °C for at least overnight and up to several months. Specimens were then rehydrated on ice through 75%, 50%, and 25% MeOH in PBST to 100% PBST, followed by an additional 10 min PBST wash at room temperature before delipidation. The delipidation solution contained 10% (v/v) THEED (Sigma Aldrich, cat. no. 87600), 5% (v/v) Triton X 100, and 25% (w/v) urea in water(42). Samples were incubated at 37 °C with shaking in delipidation solution for 30 min (developmental samples and small blastemas) or 1 h (larger limb samples), then washed three times in PBS. After a brief rinse in probe hybridization buffer (Molecular Instruments), tissues were equilibrated in hybridization buffer for 30 min at 37 °C and incubated overnight at 37 °C in hybridization solution containing 2–4 pmol HCR v3.0 probe sets (36–72 probes per gene target, SI *Appendix*, Dataset S3). Post hybridization washes comprised three rinses in 37 °C probe wash buffer (Molecular Instruments) followed by three washes in 5× SSCT. Samples received a brief room temperature rinse in amplification buffer, then were incubated overnight with 30 pmol Alexa conjugated hairpins in amplification solution prepared according to the manufacturer’s directions (Molecular Instruments). Finally, tissues were washed four times in 5× SSCT and mounted in a sucrose based mounting medium for imaging on a Zeiss 880 with Airyscan.

### Comparative genomic analysis

Predictions of translated protein sequences were obtained from putative full-length axolotl genes denoted Sp6 (AMEX60DD201010209.1) and Parpi_0005297/Sp8 (AMEX60DDU001006136.1) via UCSC Genome Browser (https://genome.axolotl-omics.org) with the March 2020 (ambMex 6.0-DD) assembly. Orthologous SP6 and SP8 protein sequences were aligned with COBALT (NCBI Constraint-Based Multiple Alignment Tool, web interface) using default settings— composition-based statistics and low-complexity filtering enabled, with automatic constraint collection from conserved domains and protein profiles—and residue conservation was visualized with the tool’s built-in conservation coloring. Phylogeny was inferred from this alignment in MEGA X (Kumar et al., 2018), and the tree was drawn to scale with branch lengths expressed as substitutions per site. For 3D visualization, homology models of each species’ SP6 and SP8 were generated in SWISS-MODEL; templates were selected automatically based on highest GMQE, sequence identity, and coverage, and the models with the best QMEAN and stereochemical assessments were retained, with no additional manual refinement or energy minimization beyond SWISS-MODEL defaults.

### Single-cell RNA sequencing

Dissociated cells from zebrafish caudal fin were prepared by the method of Thompson et al. 2020(43). Regenerating caudal fin samples were collected at time points ranging from 1 to 4 dpa via excision of the distal most 2 mm with a razor blade. For uninjured controls (0 dpa), a 2 mm section of mid caudal fin was excised that corresponded to the sampled area in injured fish. Cells were dissociated with 1.18 U/ml Liberase DH (Roche) in Hank’s Balanced Salt Solution (Gibco) with stirring at 37 °C for 2 h, collected in 15 min increments, then strained through a 40 μm nylon cell strainer (Corning). Cell suspensions were stained with SYTOX Blue (Invitrogen) and sorted for exclusion of the dye via FACS (Astrios Cell Sorter via Duke Cancer Institute Flow Cytometry Core Facility). In experiments examining basal epidermal cell populations, cells were further sorted by endogenous red fluorescence of the *krt1 19e:h2az2a mCherry* transgene and exclusion of green fluorescence from the *krt4:LY EGFP* transgene, which mark basal epidermal and superficial epidermal populations, respectively. FACS-sorted suspensions were then resuspended in PBS with 400 μg/mL non-acetylated bovine serum albumin (Sigma). An aliquot of the suspension was counted by hemacytometer and checked for viability with 0.4% trypan blue (Sigma), which ranged from 60-98% between samples.

Cells were encapsulated in gel bead emulsions and barcoded using the Chromium Controller (10x Genomics) with target yields of 6,000-10,000 cells. Preparation of cDNA and Illumina sequencing libraries was carried out according to the Chromium Single Cell 3’ Reagents Kit v.2 (10x Genomics). Libraries were sequenced with either the NextSeq 500 (Duke Center for Genomic and Computational Biology, NC) or DNBseq (BGI Americas Corporation, MA).

CellRanger (v 7.0.0) was used to process raw sequencing data before subsequent analyses with danRer11 reference data. CellBender (v 0.3.0) was used to create the filtered count matrix with parameter ’--total-droplets-included 10000 --fpr 0.05 --epochs 150’ and expected cells 5000 for injured, 2 dpa, 3 dpa, 6000 for 1dpa and 4000 for uninjured samples. The Seurat objects for downstream analysis and visualizations were created from the output files of the pipeline by collecting the filtered count matrix, clusters and UMAP reductions. The doublets were removed by package DoubletFinder (v 2.0.6) and filtered by ’nFeature_RNA > 200 & nFeature_RNA < 4000 & percent.mt < 25’. Combined data were integrated by the anchors from top 20 PCAs (principal components). The batch effects were removed by harmony package (v 1.2.3). The top 18 PCAs were used to define the cluster with resolution 0.3. The top 12 PCAs were used to re-cluster the Basal Epidermal Keratinocytes. Differential gene expression analysis was conducted by Seurat (v 5.3.0) “FindMarkers” function using “Wilcoxon rank-sum test” as the test method. RNA velocity were analyzed by velocyto (v 0.17.17). The continuous gene expression topologies were visualized by SPRING-dev (commit efe6ab4). The distribution of pseudotime by samples were performed by URD R package(v 1.1.1).

### Micro-computed tomography

Harvested digit samples were fixed in 4% paraformaldehyde overnight at 4 degrees C, dehydrated with 70% ethanol for 3-4 hours at room temperature, and air-dried for several days at room temperature. Imaging was conducted using a Nikon XTH 225 ST computed tomography scanner at the Duke Shared Materials Instrumentation Facility using the following conditions: 150 kV beam energy, beam current 51 μA, isotropic voxel size of 7 μm, power 7.6 W, exposure 708 ms, gain 24 dB, tilt 0 degrees, and 2 frames per projection. Raw image data were reconstructed using Nikon CT Pro 3D software and imported into FIJI(44) for further analysis. Threshold values for each image were set using the maximal value at which background pixelation disappeared.

Each phalangeal region of interest (ROI) was then isolated. The total bone length was calculated from the midpoint of the joint face to the distal tip using the Multipoint tool. The regenerated woven bone was then computationally isolated and regenerate length was calculated using the midpoint of the proximal face of the ROI and the distal tip. Both the whole bone and regenerate bone ROIs were analyzed with the BoneJ plug-in(45) with the Analyze Particle tool to calculate the total bone volume and with the Trabecular Thickness function to calculate the average trabecular thickness for each ROI. Analysis of micro-CT data was conducted in a blinded manner by two technicians who did not perform the scanning nor reconstruction. *P-*values were calculated by ANOVA using GraphPad Prism 10.

### Bulk RNA sequencing

Regenerating mouse digit tips were harvested and trimmed to exclude all tissue proximal to the distal interphalangeal joint. Digits were flash frozen in a mortar immersed in liquid nitrogen, then pulverized with a pestle. Powderized samples were then transferred to a Dounce homogenizer with Trizol (Invitrogen, CA) for an additional stage of homogenization. RNA was extracted from the Trizol mixture according to the manufacturer’s instructions, followed by clean up with an RNA Clean & Concentrator Kit (Zymo Research, CA).

Generation of RNA libraries and sequencing was performed by Innomics Inc (San Jose, CA). RNA samples were first enriched for mRNA containing poly-A tails using oligo(dT) beads, followed by fragmentation. Fragmented mRNA molecules were reverse transcribed into first-strand cDNA using random primers. Second-strand cDNA was synthesized using dUTP instead of dTTP to facilitate strand-specific sequencing. The resulting cDNA was end-repaired and 3’ adenylated, followed by the ligation of adaptors to the ends of the 3’ adenylated cDNA fragments. PCR amplification was then performed to enrich the cDNA library, which underwent quality control assessments. Libraries were circularized and amplified to form DNA nanoballs (DNBs).

DNBs were loaded onto the DNBSEQ-T7 sequencing platform according to the manufacturer’s protocol. High-throughput sequencing was performed, generating paired-end reads of 150 bp in length. The raw sequencing reads were QCed and filtered with SOPAnuke(46) to remove low quality reads. Final sequencing quality was assessed using FastQC.

Analysis was conducted at the Duke Regeneromics Core. RNA-seq reads were trimmed by Trim Galore (v0.6.4) and mapped with STAR(47) v2.6.1d, with parameters --twopassMode Basic -- runDirPerm All_RWX and supplying the Ensembl GRCm38 annotation) to the mouse genome (GRCm38). The mapped reads were counted using featureCounts (48) v1.6.4. Bioconductor package DESeq2(49) (v1.28.1) was employed to analyze differential expressions (DE) with litter and genotype information. Gene Ontology and KEGG enrichment tests were performed to analyze enriched biological processes by clusterProfiler(50) v3.16.1. The volcano plots were created by EnhancedVolcano (v1.6.0). The coverage depth was normalized by deeptools(51) v3.1.3 using RPKM for RNA-seq. TPM values were quantified from Salmon(52) v1.2.1 quantification and summarized via tximport v1.16.1(53). Differentially expressed genes were defined at p < 0.05 and log₂ fold change > 0.58. Gene ontology/pathway enrichment was performed with Metascape(54) . Statistically enriched terms (GO/KEGG, canonical pathways, Hallmark gene sets) were identified using cumulative hypergeometric testing; p values and enrichment factors were used for filtering. Significant terms were hierarchically clustered by Kappa similarity, and clusters were defined at a Kappa score threshold of 0.3.

### AAV production and administration

Recombinant AAV vectors were prepared using suspension HEK293 cell cultures as previously described(28, 55). Suspension cells were triple-plasmid-transfected with PEI, containing pITR (0.3 ug/mL), pXX680 (0.6 ug/mL), and RepCap (0.5 ug/mL). Media was harvested at 6 days post-transfection with cells pelleted and discarded by centrifugation. Media was incubated with 12% polyethylene glycol (PEG) overnight at 4 °C. Media was then centrifuged at 4,000 x G at 4 °C for 45 minutes to precipitate PEG containing AAV. PEG was resuspended in AAV formulation buffer, containing 1 mM MgCl2 and 0.001% F-68 in dPBS. Resuspended PEG and AAV was DNase treated at 37 °C for 1 hour, followed by iodixanol gradient purification via ultracentrifugation at 30,000 RPM at 17 °C overnight. Iodixanol gradients consisted of 60%, 40%, 25%, and 17% densities. Following centrifugation, 550 ul fractions were collected starting from the 25%-40% border and ending at the 40%-60% border. All fractions were titered for AAV by qPCR using primers targeting the ITR region. Iodixanol fractions containing the highest titer were selected for downstream desalting and buffer exchange. Desalting was performed using manufacturer’s instructions with a PD MidiTrap G-25 column (Cytiva). Following desalting, buffer exchange with sterile AAV formulation buffer was performed according to the manufacturer’s instructions with a Pierce high-capacity endotoxin removal spin column (Thermo Scientific). Purified final AAV titer was measured by QIAcuity Digital PCR (dPCR) system with QIAcuity probes targeting AAV2-ITR (Qiagen, #250102). Titered AAV was systemically delivered in 2-6-month-old mice via retro-orbital injection at a dosage of 2.5x10^13^ viral genomes/kg body weight.

### Bioluminescence imaging

Mice were anesthetized as described above for digit amputations and maintained on a 37 °C heated stage; ophthalmic ointment was applied to prevent corneal drying. D-luciferin (Promega, Madison, WI) was prepared immediately before use per the manufacturer’s instructions and administered intraperitoneally at 10 µL/g body weight. Animals were positioned consistently in the imaging chamber and imaged on an IVIS Lumina (PerkinElmer) using open emission filter, f/stop = 1, medium binning, and exposures between 1–60 s (auto exposure enabled and kept within the linear range; identical settings used for all animals). Images were acquired beginning 5 min post injection and then every 2–3 min for 15–20 min; the frame with the peak signal (typically 10–15 min post injection) was used for comparisons. Quantification was performed in Living Image software (PerkinElmer) using fixed size, background subtracted regions of interest (ROIs) placed over the target site and reported as total flux (photons/s) and/or radiance (photons/s/cm²/sr). All acquisition settings, ROI sizes, and analysis steps were held constant across animals and time points.

## Acknowledgments

The authors would like to acknowledge Veronica Han and Choiselle Marius for their assistance with histologic staining. We would like to thank the Wake Forest Biology Microscopic Imaging Core RRID:SCR_021975 for access and assistance with confocal and fluorescent stereoscope imaging.

This work was supported by grants from the National Institute of Arthritis and Musculoskeletal and Skin Diseases to D.B. (K08 AR074529), the Eunice Kennedy Shriver National Institute of Child Health and Human Development to D.B. and K.D.P. (R01 HD115266) and to K.D.P. (R01 HD105033), discretionary funds from Duke University School of Medicine and Morgridge Institute for Research to K.D.P., the Duke Office of Physician-Scientist Development Strong Start Award to D.B., and Wake Forest University institutional funding to J.C. T.C was partially supported by funding from the Wake Forest Center for Molecular Signaling. K.K. was funded by the Deutsche Forschungsgemeinschaft (DFG, German Research Foundation – 547486810). L.S. was supported by the Eunice Kennedy Shriver National Institute of Child Health and Human Development (F32 HD103376). A.A. was funded in part by the National Institutes of Health (R01HL089221, R01DK134408, U01AI170064). A.A. is a co-founder and director at Torque Bio. A.A. is listed as an inventor on patents covered in the subject matter of this manuscript.

This work was performed in part at the Duke University Shared Materials Instrumentation Facility (SMIF) (RRID:SCR_027480), a member of the North Carolina Research Triangle Nanotechnology Network (RTNN), which is supported by the National Science Foundation (award number ECCS-2025064) as part of the National Nanotechnology Coordinated Infrastructure (NNCI).

Figure schematics were created with BioRender.com.

---

**Fig. S1.**
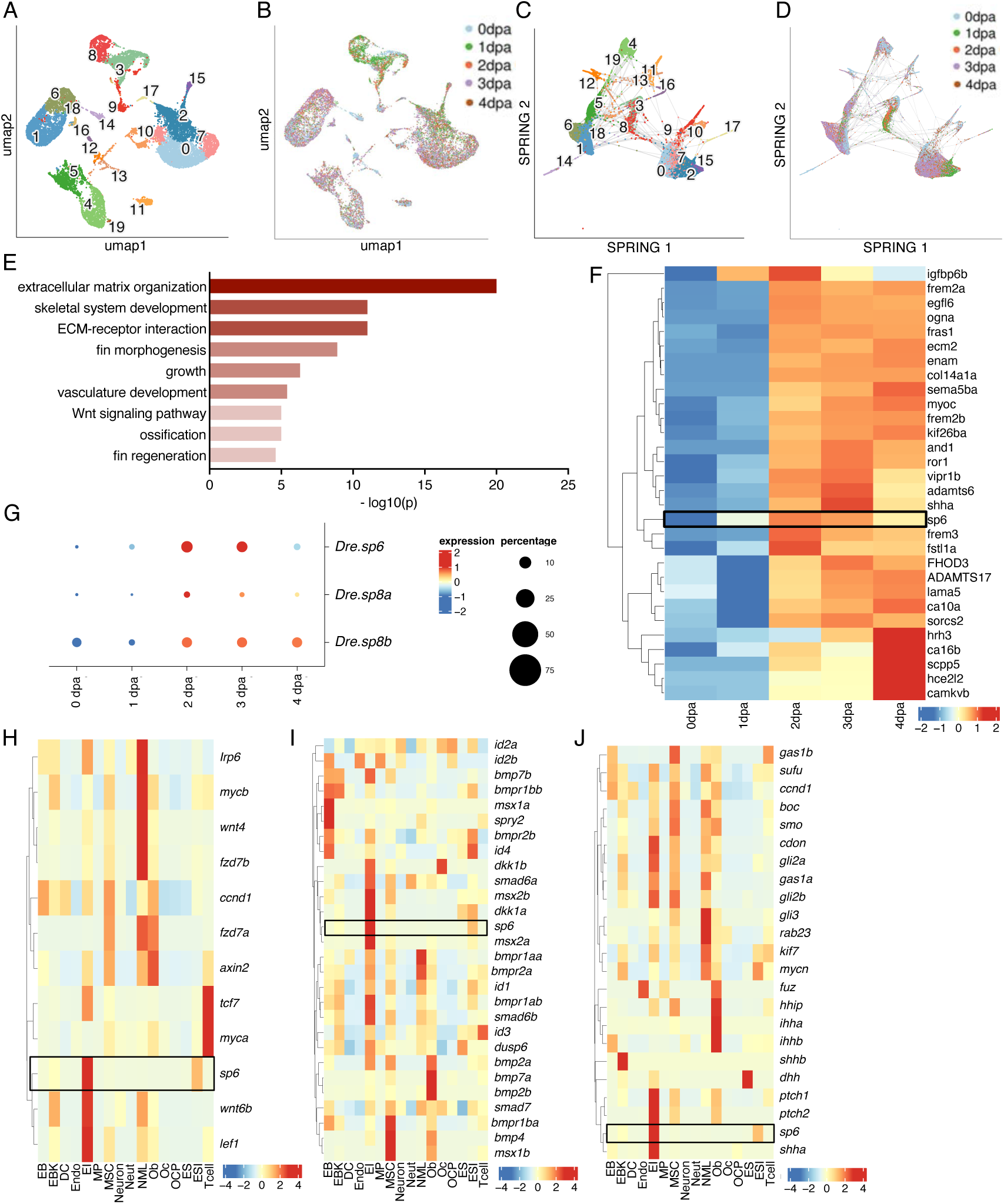
scRNA-seq identifies an *sp6/sp8*-high intermediate epithelial compartment in regenerating fins. (A) UMAP of the integrated scRNA-seq dataset showing clusters 0–19. (B) The same UMAP colored by regeneration stage (0–4 dpa). (C–D) SPRING layouts of the integrated dataset colored by cell cluster (C) or stage (D). (E) GO Biological Process enrichment for marker genes of the intermediate epithelium (IE; cluster 6). (F) Heatmap of the top 30 IE marker genes across stages (0–4 dpa); values are scaled average expression with hierarchical clustering of genes. (G) Dot plot of *Dre.sp6*, *Dre.sp8a*, and *Dre.sp8b* across stages; color denotes mean expression (dark blue = low, red = high) and dot size indicates the percent of cells expressing the gene. (H–J) Pathway heatmaps across annotated cell types (columns as in panel B): Wnt/β-catenin (H), BMP/TGFβ (I), and Hedgehog (J). Rows list pathway components/targets (gene symbols; red labels mark canonical ligands/targets). Values are row-scaled (z-score). Black boxes highlight *sp6* enrichment.

**Fig. S2.**
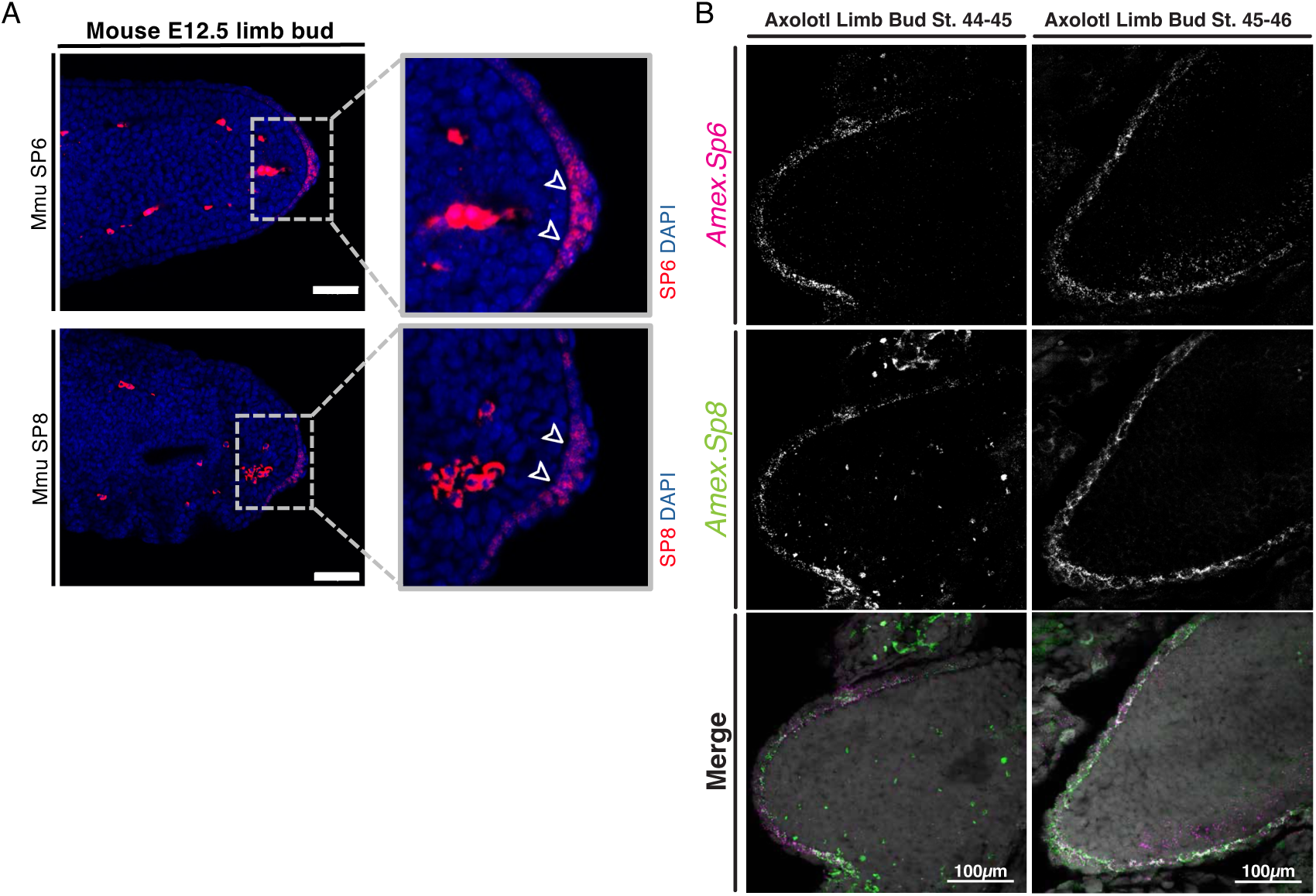
Conserved SP6/SP8 expression during embryonic limb development in mouse and axolotl. (A) Immunofluorescence for Mmu SP6 (top) and Mmu SP8 (bottom) on sagittal sections of mouse limb buds at E12.5. Signal (red) is enriched in the distal limb ectoderm, including the apical ectodermal ridge (AER; arrowheads); nuclei are counterstained with DAPI (blue). Dashed boxes indicate regions shown at higher magnification at right. Scale bars, 100 μm (overview) and 20 μm (insets). (B) Wholemount fluorescent in situ hybridization of *Amex.Sp6* and *Amex.Sp8* in axolotl limb buds at early (Stage 44-45, left) and mid-limb bud growth (Stage 45-46, right) with DAPI merge (bottom panel). Scale bars, 100 μm.

**Fig. S3.**
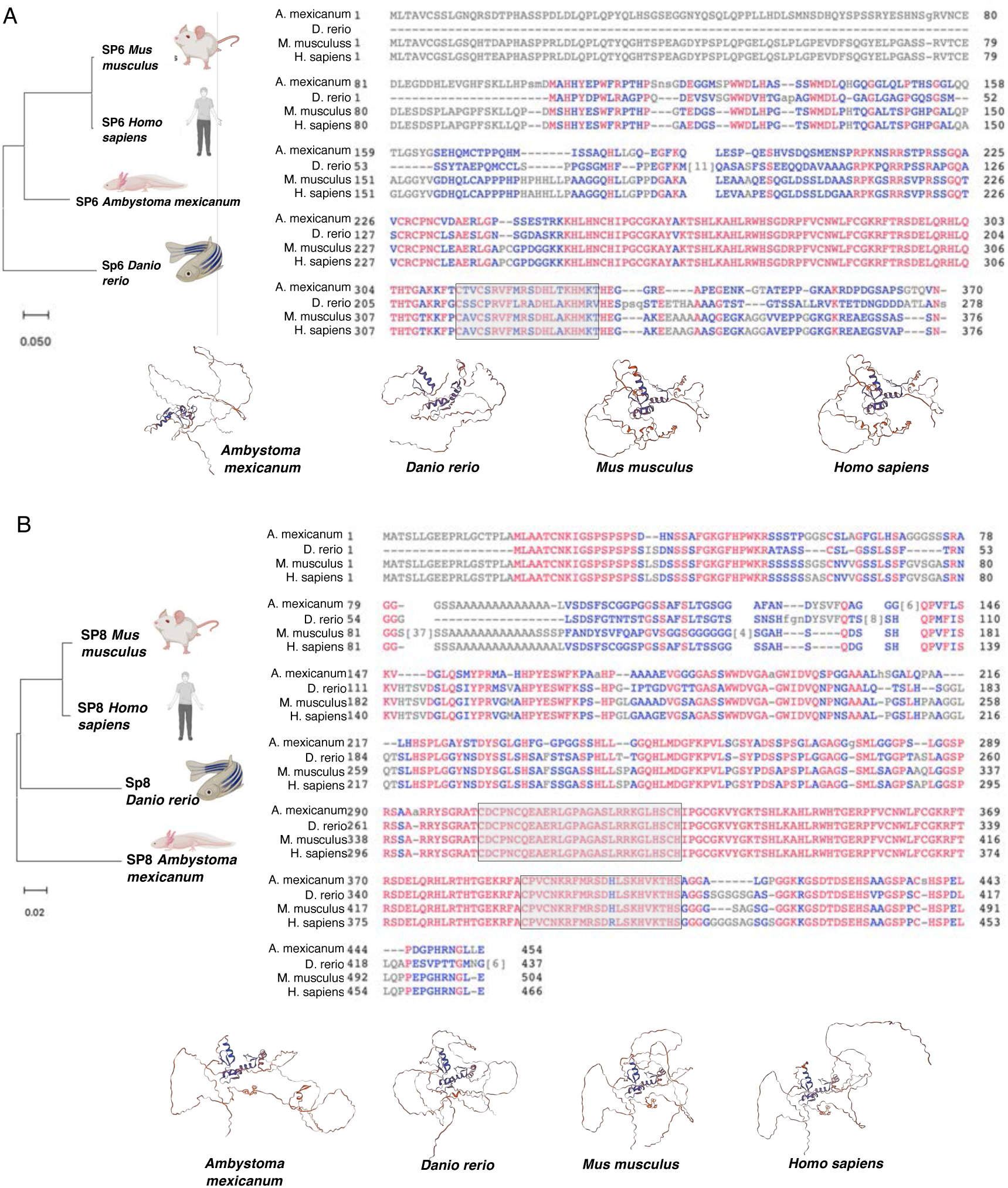
Evolutionary conservation of Sp6 and Sp8. (A) *Sp6*. Left, phylogenetic tree of SP6 orthologs from *Ambystoma mexicanum* (axolotl), *Danio rerio* (zebrafish), *Mus musculus* (mouse), and *Homo sapiens* (human); scale bar denotes substitutions per site. Right, multiple-sequence alignment of proteins; the C-terminal C2H2 zinc-finger DNA-binding array is boxed. Bottom, ribbon representations of the 3D protein structures for each species. (B) *Sp8*. As in (A), shown for SP8 orthologs from the same four species: phylogeny (left), protein alignment with the annotated C2H2 zinc-finger array (right), and ribbon representations of the 3D structures (bottom).

**Fig. S4.**
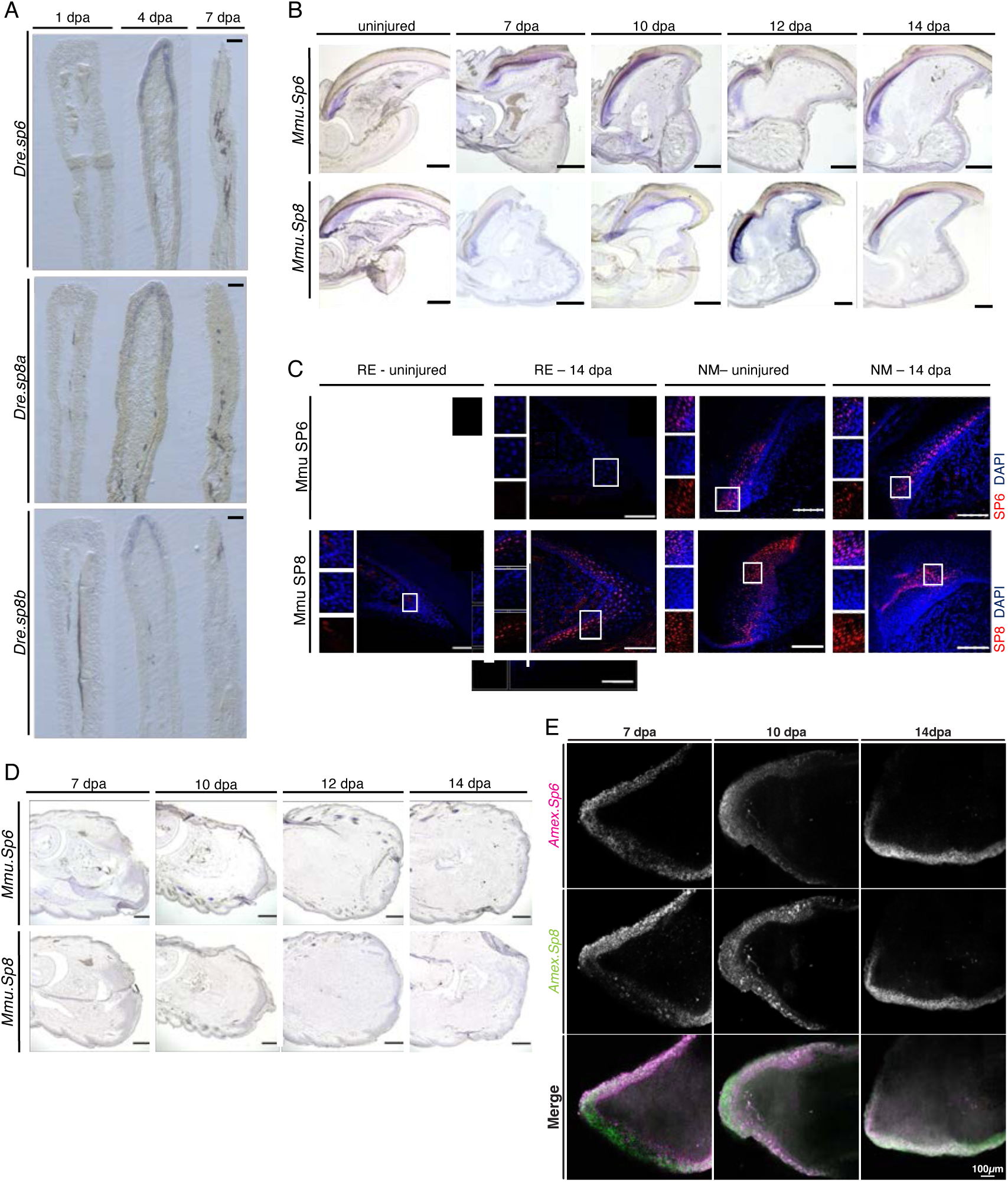
Regeneration-specific *Sp6* and *Sp8* expression across species. (A) In situ hybridization (ISH) for *Dre.sp6*, *Dre.sp8a*, and *Dre.sp8b* on longitudinal sections of regenerating caudal fins at 1, 4, and 7 days post-amputation (dpa). Signal is enriched in the distal wound/regenerate epidermis. Scale bars, 50 μm. (B) ISH for *Mmu.Sp6* (top row) and *Mmu.Sp8* (bottom row) on sagittal sections of mouse digit tips (P3 amputation), shown for uninjured and 7, 10, 12, and 14 dpa. Signal localizes to the distal nail epithelium/wound epithelium of regenerating digits. Scale bars, 200 μm. (C) Immunofluorescence for Mmu SP6 (top) and Mmu SP8 (bottom) on sagittal sections of mouse digit tips (P3 amputation), comparing nail matrix (NM) and regenerating epidermis (RE) in uninjured and 14 dpa digits. Signal (red) marks SP6/SP8; nuclei are counterstained with DAPI (blue). Boxes denote regions shown at higher magnification in the flanking insets. Scale bars, 50 μm. (D) ISH for *Mmu.Sp6* (top row) and *Mmu.Sp8* (bottom row) mRNAs on sagittal sections of P2-amputated (non-regenerating) mouse digits at 7, 10, 12, and 14 dpa. Signal is low to undetectable across time points in this injury context. Scale bars, 200 μm. (E) Wholemount fluorescent ISH of *Amex.Sp6* and *Amex.Sp8* in lower limb (mid radius/ulna) at 7, 10, 14 days post amputation (dpa) with DAPI merge (bottom panel). Scale bar, 100 μm.

**Fig. S5.**
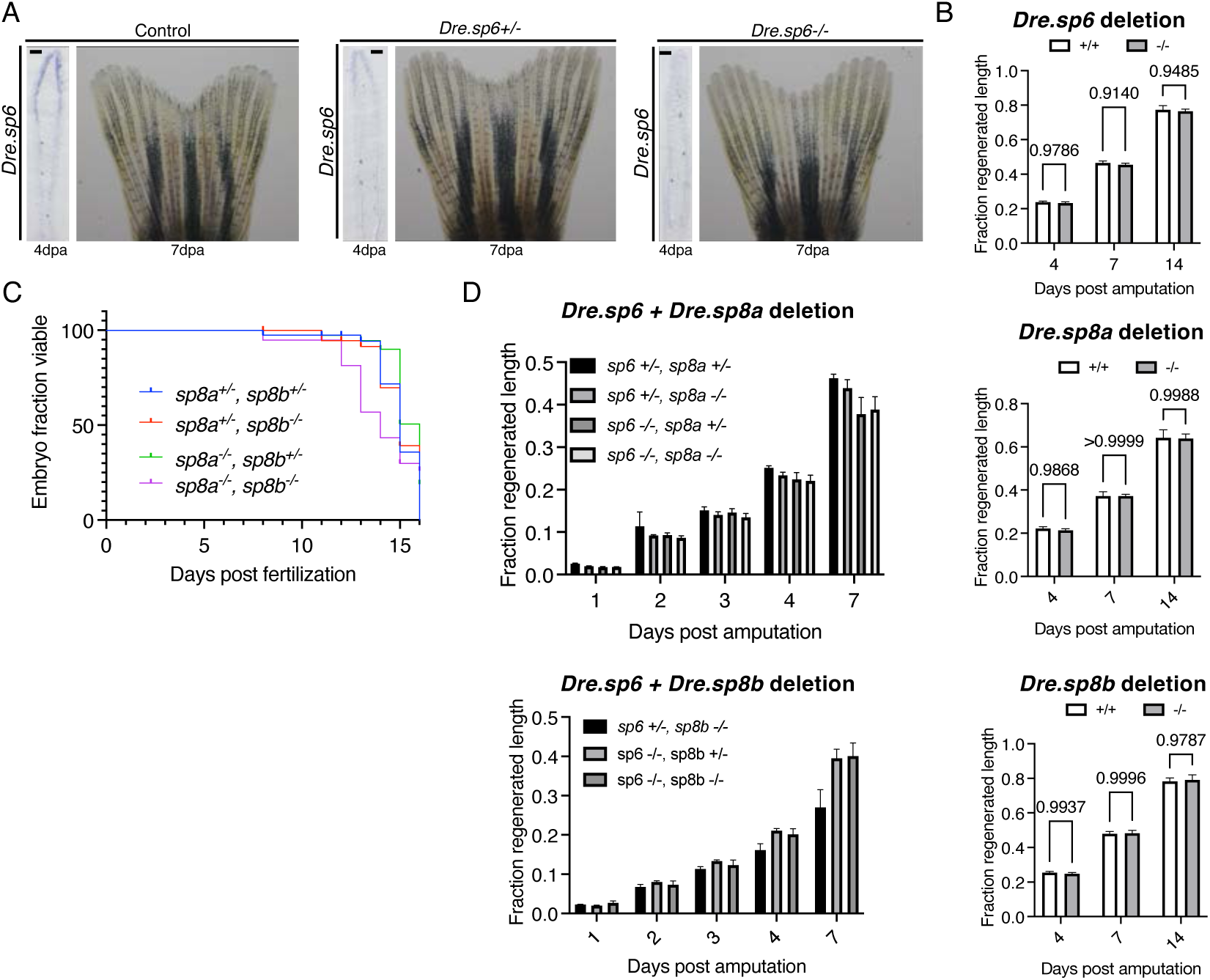
Zebrafish fin regeneration in single and compound *Sp* mutants. (A) *Dre.sp6* in situ hybridization at 4 dpa (left insets) and gross fin images at 7 dpa (right) for control, *sp6*+/–, and *sp6*–/– fish. ISH panels show endogenous *Dre.Sp6* mRNA in the regenerating fin tip. Scale bar, 50 μm (ISH panels). (B) Quantification for single-gene deletions (Dre.*sp6*, Dre.*sp8a*, Dre.*sp8b*). Fraction regenerated length (injured fin normalized to the contralateral uninjured fin) at 4, 7, and 14 dpa. Data are mean ± SEM; *n* = 8–10 fish per group, with 4 rays tracked per fish; *p*-values are indicated above brackets. (C) Viability of zebrafish embryos carrying *Dre.sp8a/Dre.sp8b* alleles, plotted as Kaplan–Meier– style survival curves from fertilization to 15–16 days post fertilization. Genotypes: *sp*8a+/−;*sp*8b+/− (blue), *sp*8a+/−;*sp*8b−/− (red), *sp8a*−/−;*sp8b*+/− (green), and *sp8a*−/−;*sp8b*−/− (purple). (D) Early regeneration time courses (1–7 dpa) for compound genotypes combining *Dre.sp6* with *Dre.sp8a* (left) or *Dre.sp8b* (right). Fraction regenerated length plotted for the indicated allele combinations; data are mean ± SEM.

**Fig. S6.**
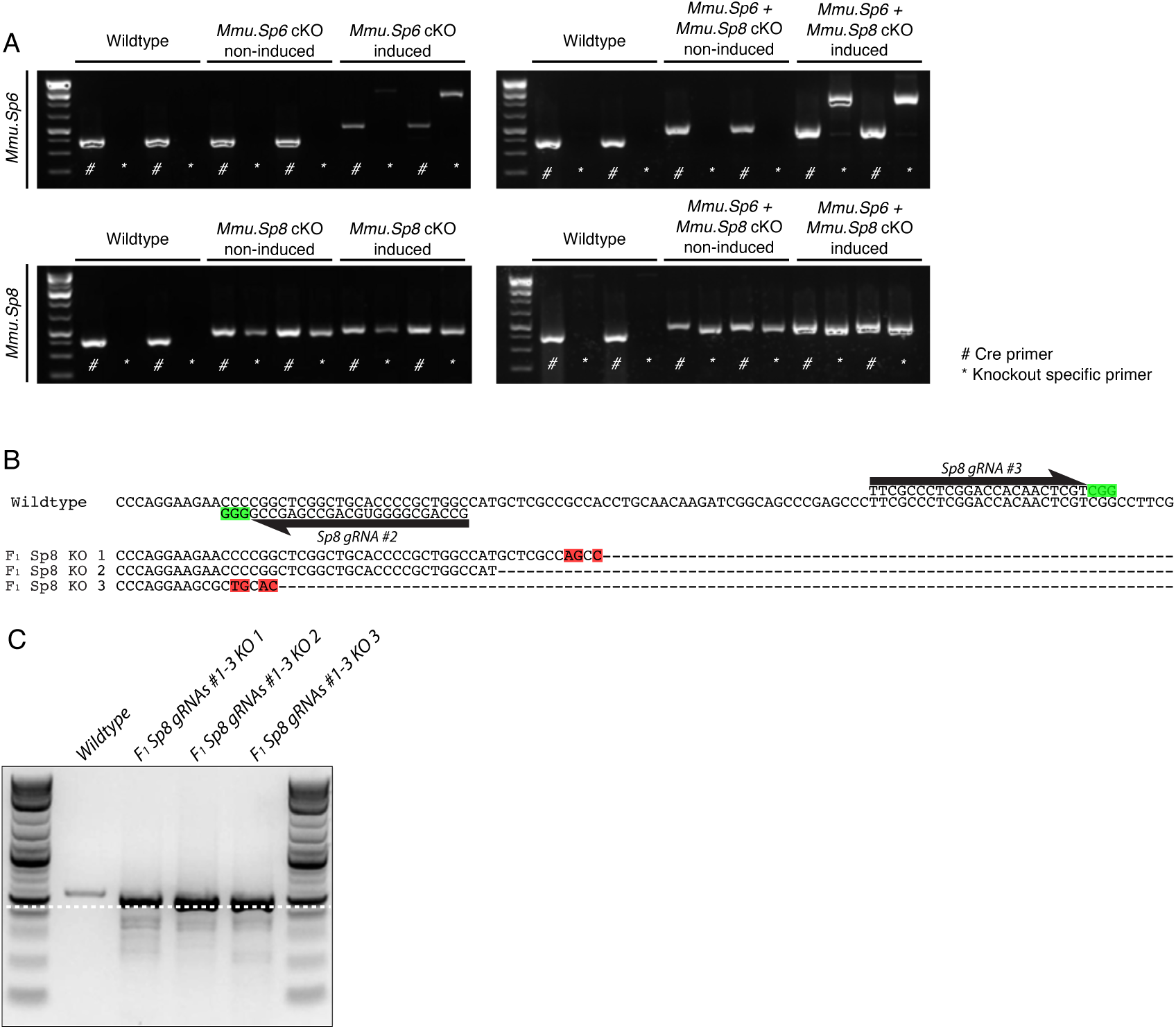
Genotyping of Sp knockouts in axolotl and mice. (A) Agarose gel electrophoresis of genomic PCR products of conditional deletion of *Mmu.Sp6* (top) and *Mmu.Sp8* (bottom). Lanes show wild type, single cKO (non-induced vs. induced), and double cKO (non-induced vs. induced) as labeled. “#” denotes the Cre transgene amplicon; “*” denotes the knockout–specific amplicon. DNA ladder at left. (B) Consensus sequence from genomic PCR NGS reads covering the wildtype *Amex.Sp8* locus within a region including complementary sites for *Amex.Sp8* gRNA #2 and *Amex.Sp8* gRNA #3. The gRNA protospacer adjacent motif (PAM) sequence is indicated in green. Genomic reads from three F1 *Amex.*Sp8 gRNAs #1-3 hemizygous individuals show truncations and mutations (red) between gRNA #2 and gRNA #3. (C) Agarose gel electrophoresis of genomic PCR products of wildtype and hemizygous knockout individuals sequenced in (A). Dashed line indicates approximately 500bp DNA fragment size.

**Fig. S7.**
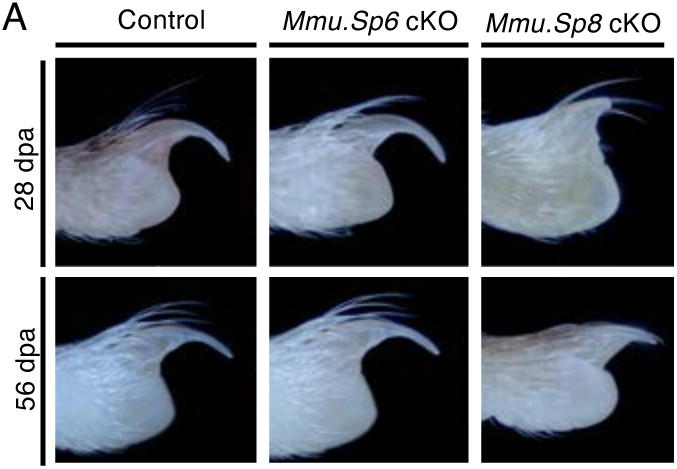
Bone regeneration and gross morphology in epithelial *Mmu.Sp* knockouts. (A) Representative whole-mount images at 28 and 56 dpa comparing nail growth and distal digit morphology across control, *Mmu.Sp6* cKO, and *Mmu.Sp8* cKO groups.

**Fig. S8.**
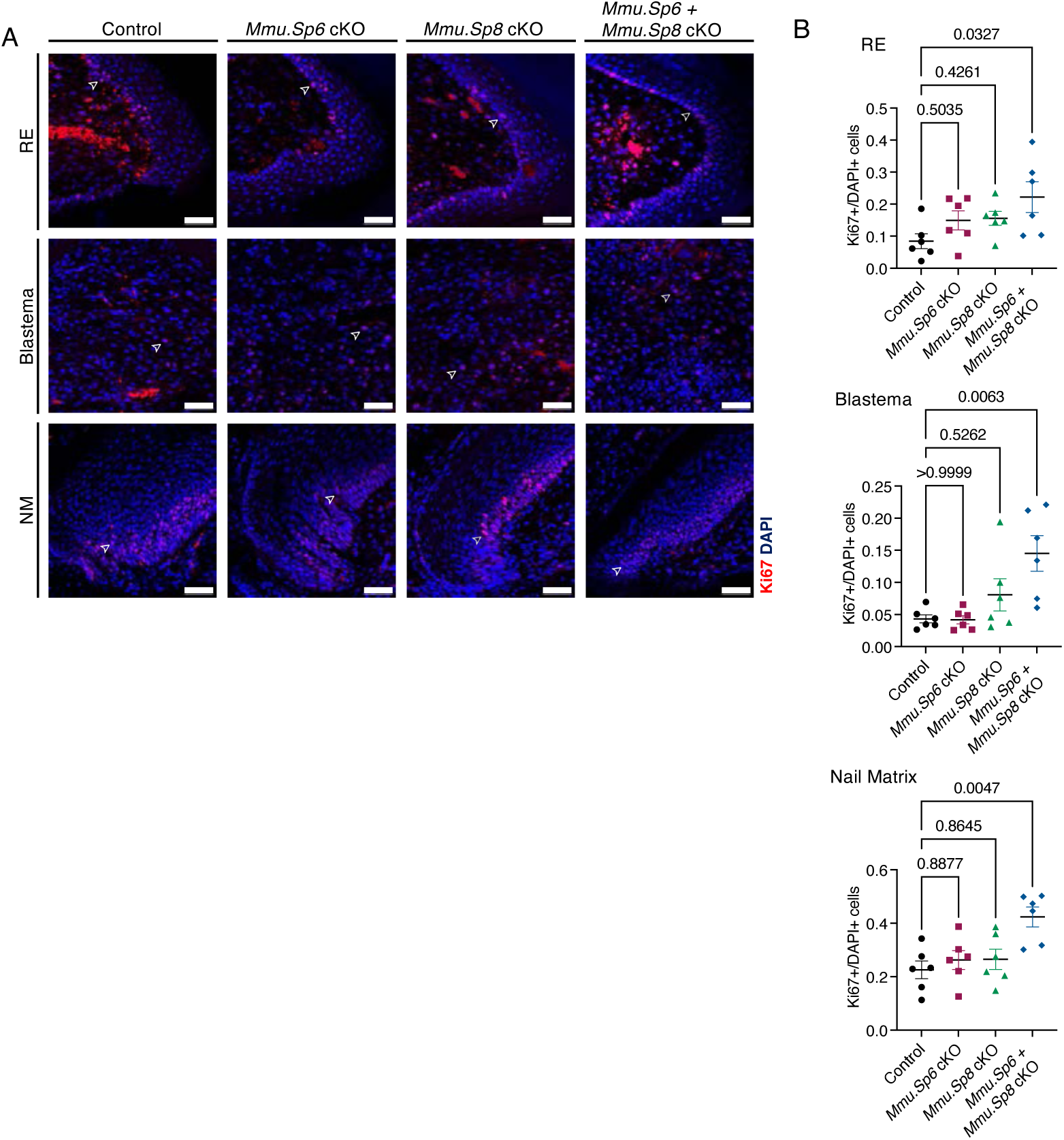
Epithelial Sp6/Sp8 deletion alters proliferation during digit regeneration. (A) Immunofluorescence for Ki67 (red) in sagittal sections of mouse digit tips (P3) from control, *Mmu.Sp6* cKO, *Mmu.Sp8* cKO, and Mmu.*Sp6* + *Mmu.Sp8* cKO animals. Images are shown for the regenerating epidermis (RE), blastema, and nail matrix (NM) at 14 dpa. Nuclei are counterstained with DAPI (blue); arrowheads mark Ki67⁺ cells. Scale bars, 50 μm. (B) Quantification of proliferation as the fraction of Ki67⁺/DAPI⁺ cells in RE, blastema, and nail matrix at 14 dpa (*n* = 6 per group). Points represent individual digits; bars indicate mean ± SEM. *p*-values for the indicated comparisons are shown.

**Fig. S9.**
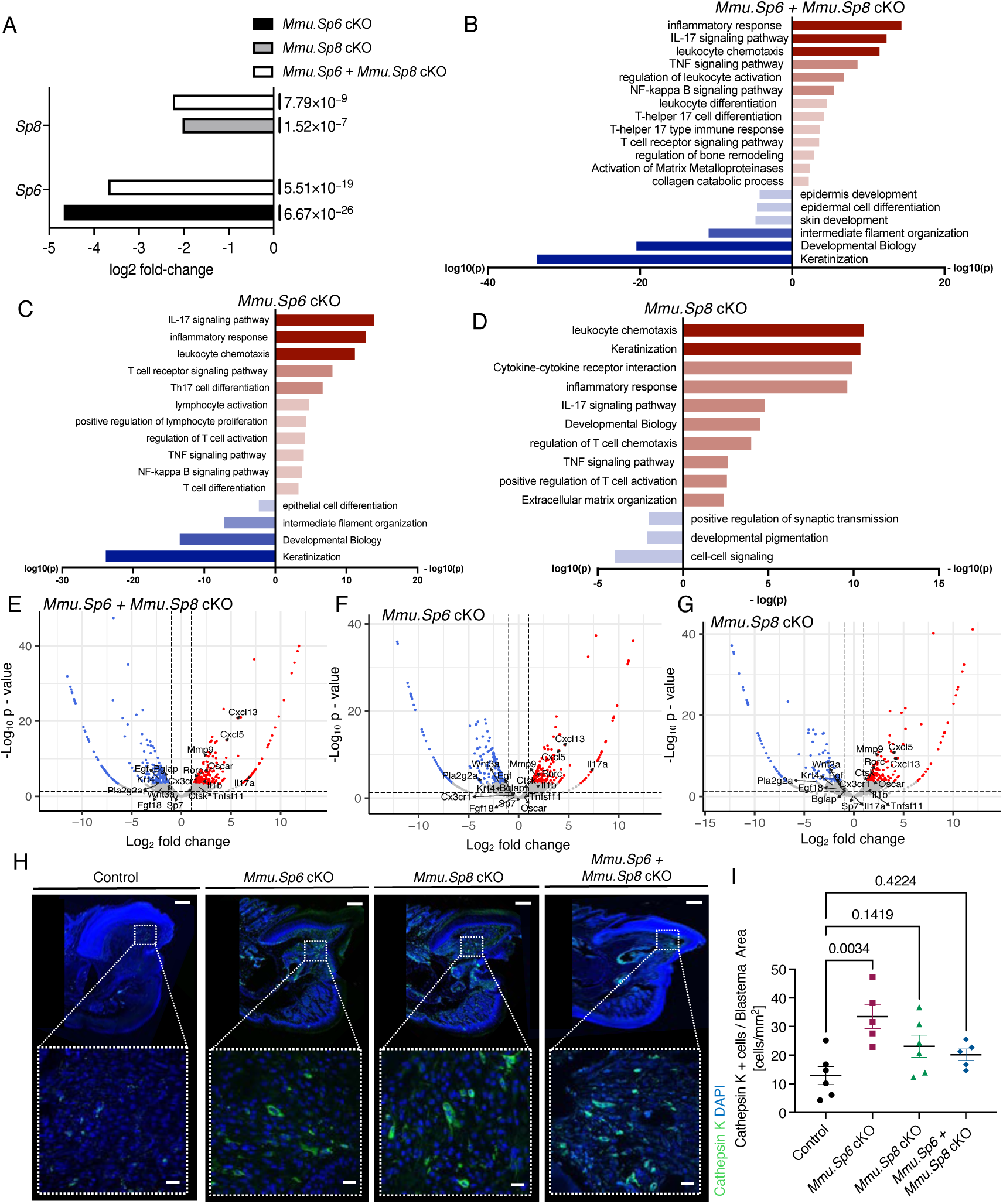
Bulk RNA-seq of regenerating digits following epithelial Sp6 or Sp8 deletion. (A) Horizontal bar plot showing log₂ fold-change in expression (relative to control) for *Sp6* and *Sp8* in *Mmu.Sp6* cKO, *Mmu.Sp8* cKO and Mmu.*Sp6* + *Mmu.Sp8* cKO groups. Bars denote genotype: black, Sp6 cKO; gray, Sp8 cKO; open, Sp6 + Sp8 double cKO. Numbers to the right of each bar are the corresponding p values. *n* = 3 biological replicates per group. (B) Gene Ontology (GO) analysis of bulk RNA-seq at 14 dpa, showing pathways upregulated (red) or downregulated (blue) in *Sp6+Sp8* cKO digits relative to controls. (C) Gene Ontology (GO) enrichment of DE genes at 14 dpa, showing pathways upregulated (red) and downregulated (blue) in *Sp6*^Krt14-KO^ relative to controls. (D) GO enrichment of DE genes at 14 dpa, with pathways upregulated (red) and downregulated (blue) in *Mmu.Sp8* cKO relative to controls. (E) Volcano plot of bulk RNA-seq data at 14 dpa comparing *Sp6+Sp8* cKO and control digits. Highlighted genes include regulators of *IL-17* signaling and inflammation (*PLA2G2A, IL1B, CX3CR1, Cxcl13, Cxcl5*), osteoclast differentiation and bone resorption (*MMP9, Oscar, Tnfsf11/RANKL, Ctsk*), osteoblast differentiation and mineralization (*Bglap, WNT3A, Fgf18*), and epidermal proliferation and wound healing (*EGF*). (F) Volcano plot for *Mmu.Sp6* cKO versus control. Highlighted genes include regulators of IL-17 signaling/inflammation (*PLA2G2A, IL1B, CX3CR1, Cxcl13, Cxcl5*), osteoclast differentiation/bone resorption (*MMP9, Oscar, Tnfsf11/RANKL, Ctsk*), osteoblast differentiation/mineralization (*Bglap, Wnt3a/Wnt4, Fgf18*), and epidermal proliferation/wound healing (*EGF*). (G) Volcano plot for *Sp8*^Krt14-KO^ versus control, highlighting the same functional categories as in (E), including IL-17/inflammatory genes (*PLA2G2A, Cxcl13, Cxcl5*), osteoclast markers/regulators (*MMP9, Oscar, Tnfsf11/RANKL, Ctsk*), and osteoblast/epidermal programs (*Bglap, Wnt4, Fgf18*). (H) Representative immunofluorescence images of Cathepsin K staining in control, Sp6 cKO, Sp8 cKO, and Sp6+Sp8 cKO digits. Scale bars: 200 μm (top), 20 μm (bottom). (I) Quantification of Cathepsin K⁺ cells in the blastema region across control, Sp6 cKO, Sp8 cKO, and Sp6+Sp8 cKO digits. Data are mean ± SEM (control and Sp8 cKO, n = 6; Sp6 cKO and Sp6+Sp8 cKO, n = 5); p-values indicated in the graph.

**Fig. S10.**
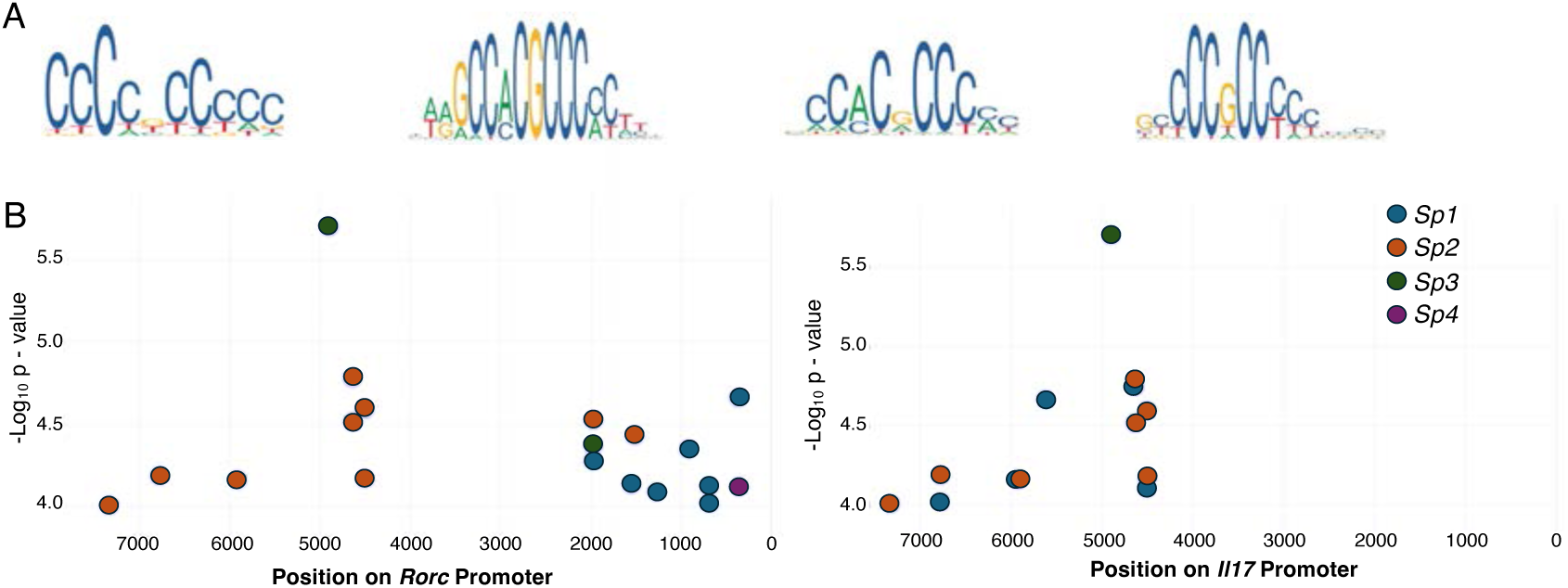
*Sp* DNA binding motifs analysis. (A) DNA binding motifs for SP1, SP2, SP3, and SP4 transcription factors illustrating consensus GC-rich binding sites recognized by each factor. (B) In silico prediction of transcription factor binding sites for SP family members (SP1–SP4) in the promoters of *Rorc* (left) and *Il17* (right). Each dot represents a predicted binding site, plotted by genomic from the transcriptional start site (TSS) and statistical significance.

**Fig. S11.**
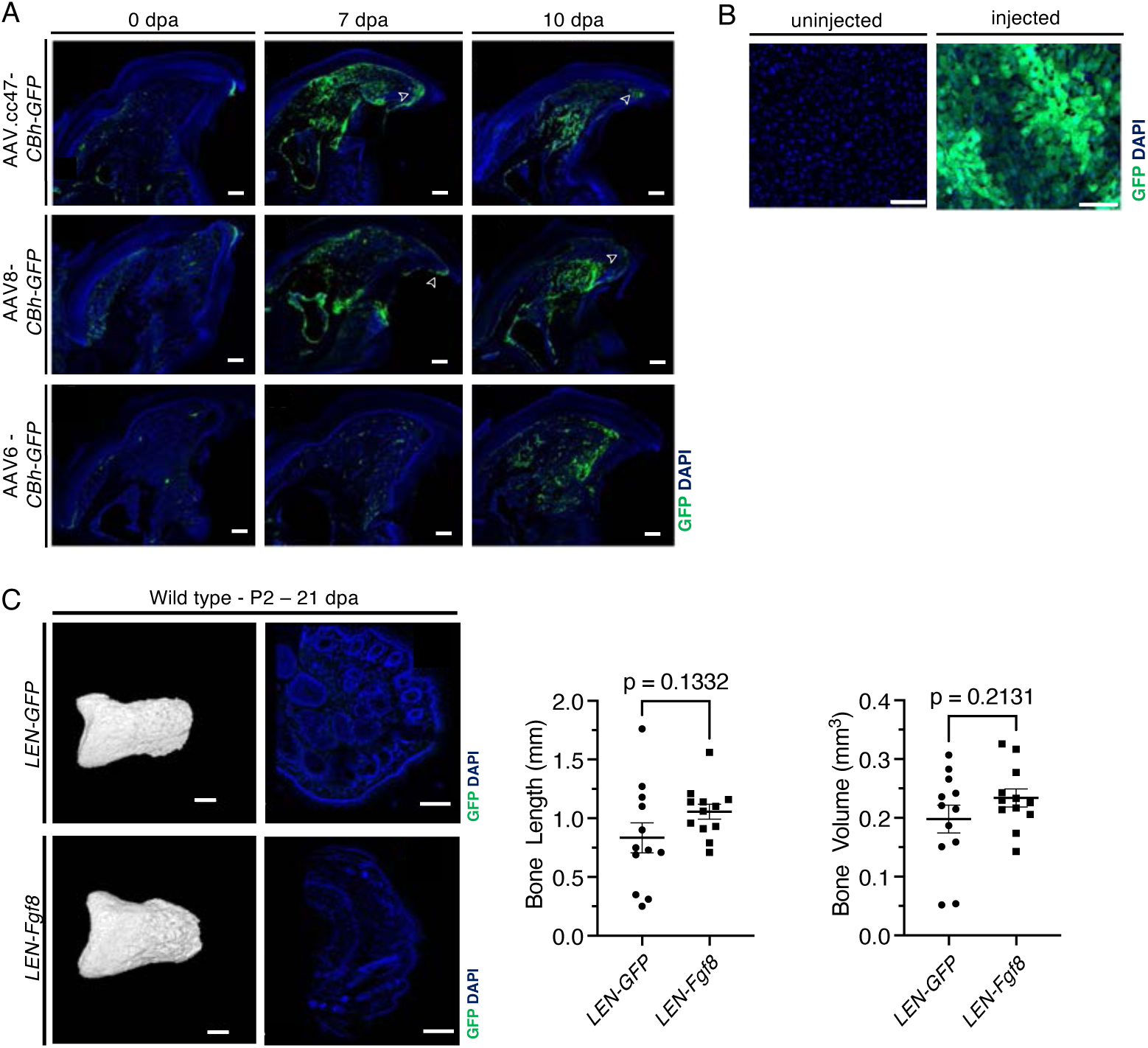
AAV-mediated gene delivery and its effects on digit regeneration outcomes. (A) Comparison of GFP immunofluorescence in P3-regenerating digit tips at 14 dpa after AAV administration at 0, 10, or 14 dpa. Different AAV serotypes (AAV.cc47, AAV8, and AAV6) carrying the *CBh-GFP* reporter were injected, and GFP signal (green) indicates transgene expression. Nuclei are counterstained with DAPI (blue). Arrows indicate GFP expression in the distal epidermis. Scale bars, 100 μm. (B) Immunofluorescence images of control (uninjected) and experimental (injected) liver samples. Robust GFP signal (green) is detected only in the injected animal, indicating successful AAV delivery, whereas the uninjected control shows background levels. Nuclei are counterstained with DAPI (blue). Scale bars, 50 μm. (C) Micro-CT reconstructions and quantitative analysis of digit regeneration in wild-type mice (P2) at 21 dpa following injection of *LEN-GFP* or *LEN-Fgf8*. Representative 3D reconstructions (left) show bone morphology and density. Immunofluorescence comparison indicates GFP expression (middle). Quantification (right) includes total bone length and bone volume. Data are presented as mean ± SEM (n = 12 per group); *p*-values are indicated. Scale bars, 200 μm.

**Table S1:**
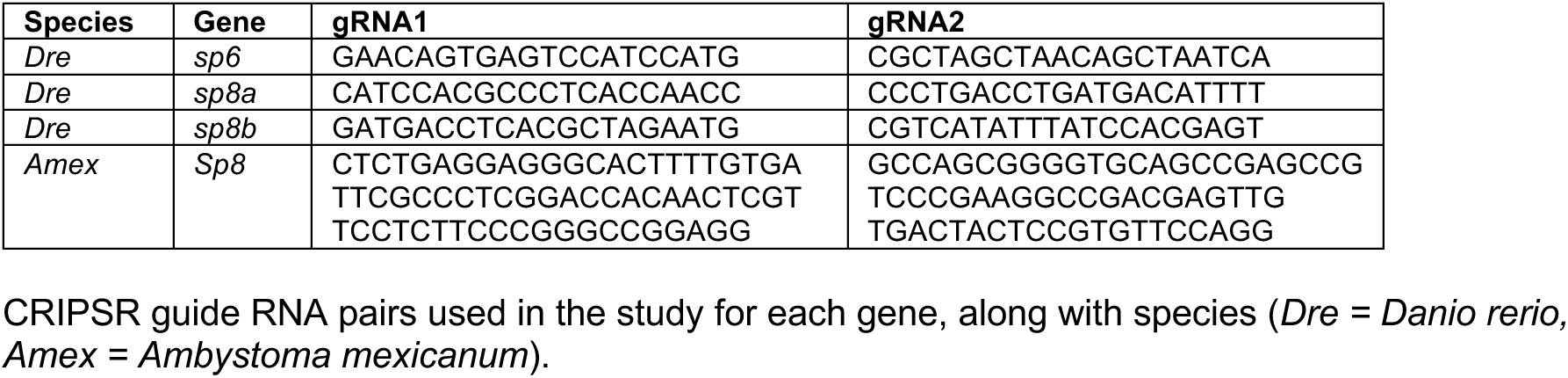
CRISPR guide RNA sequences.

**Table S2:**
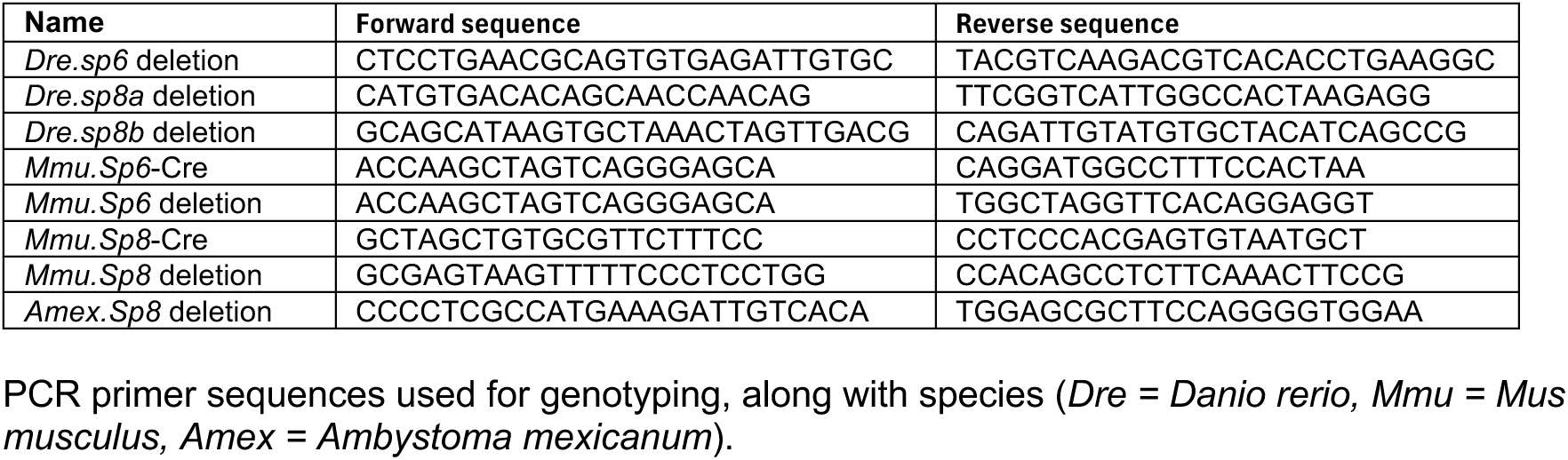
Genotyping primers.

**Other supporting materials for this manuscript (separate files):**

**Movie S1. Live imaging of *Tg(krt1*-*19e:h2az2a*-*mCherry)*; *Tg(krt4:LY*-*EGFP)* zebrafish following caudal fin amputation.**

**Dataset S1. scRNA-seq differential gene expression analysis Dataset**

**S2. Riboprobe sequences for in-situ hybridization**

